# Altered projection-specific synaptic remodeling and its modification by oxytocin in an idiopathic autism marmoset model

**DOI:** 10.1101/2022.08.24.505057

**Authors:** Jun Noguchi, Satoshi Watanabe, Tomofumi Oga, Risa Isoda, Keiko Nakagaki, Kazuhisa Sakai, Kayo Sumida, Kohei Hoshino, Koichi Saito, Izuru Miyawaki, Eriko Sugano, Hiroshi Tomita, Hiroaki Mizukami, Akiya Watakabe, Tetsuo Yamamori, Noritaka Ichinohe

**Author notes:** Correspondence should be addressed to J.N. or N.I.

## Abstract

Alterations in the experience-dependent and autonomous elaboration of neural circuits are assumed to underlie autism spectrum disorder (ASD), though it is unclear what synaptic traits are responsible. Here, we used a valproic acid-induced ASD marmoset model, which shares common molecular features with idiopathic ASD, to investigate the structural dynamics of tuft dendrites of upper-layer pyramidal neurons and adjacent axons in the dorsomedial prefrontal cortex using two-photon microscopy. In model marmosets, dendritic spine turnover was upregulated, and spines were actively generated in clusters and subsequently survived more often than in control marmosets. Presynaptic boutons in local axons but not in commissural long-range axons showed hyperdynamic turnover in model marmosets, suggesting alterations in projection-specific plasticity. Intriguingly, nasal administration of oxytocin reduced the clustered spine emergence. Enhanced clustered spine generation, possibly unique to certain presynaptic partners, may be associated with ASD and may be a potential therapeutic target.

## Introduction

Autism spectrum disorder (ASD) is a developmental disorder characterized by impairments in social communication, social interaction, and stereotyped behaviors ^1, 2^. Individuals with ASD often have learning disabilities and have difficulty learning to recognize verbal or non-verbal social information ^3^. Proper refinement of neural networks during learning is achieved by coordinated synaptic remodeling, which may be altered in ASD. Dendritic spines, which are postsynaptic protrusions that receive most excitatory inputs ^4–6^, have been observed longitudinally using two-photon microscopy in mouse models of ASD. This has made it possible to explore the spatiotemporal characteristics of synaptic remodeling in ASD. Morphological analysis of dendrites with in vivo two-photon microscopy has shown that accelerated spine generation and elimination in the motor and primary sensory cortices is a consistent phenotype in numerous ASD mouse models (BTBR, 15q11-13 duplication, Neuroligin mutant, *FMR1* knockout, *MeCP2* duplication, etc.) ^7–10^.

Clustered generation of postsynaptic dendritic spines, in which new spines form in close proximity to one another, plays a critical role in learning and memory ^11–17^. Training increases clustered spine generation, which has been found to be correlated with learning performance in corresponding brain regions ^10, 18^. Neuronal modeling studies have suggested that enhanced clustered spine generation increases memory discrimination and the storage capacity of neuronal networks ^17, 18^. Excessively clustered spines may contribute to the development of ASD symptoms, but little is known about their involvement in this condition. On the other hand, it has been reported that diversity of excitatory synapses includes several synapse types or subtypes defined by molecular and other characteristics, and that certain circuits or connectome networks prefer particular types of synapses ^19^. Changes in gene expression in ASD may alter the plasticity of specific cortical projections, which may in turn perturb the formation of neural circuits adapted to learning. However, projection-specific variations of synapse remodeling in ASD have not been explored.

The common marmoset (*Callithrix jacchus*), a small New World monkey, has attracted significant attention due to its rich repertoire of social behaviors, a well-developed prefrontal cortex (PFC) that supports high-level social ability, and gene expression networks that are similar to those in humans ^20^. In fact, marmosets are more similar to humans than rodents in terms of their synaptic proteome ^21^. We previously developed a valproic acid (VPA)-induced ASD model in the common marmoset ^22^. VPA is an antiepileptic drug and also functions as a histone deacetylase inhibitor that may epigenetically increase the risk of ASD in humans by suppressing histone deacetylases in the fetal brain. Despite the fact that VPA administration was the sole environmental factor contributing to the development of ASD in this model, and there were no genetic variations, it was sufficient to induce gene expression changes in juvenile marmosets that were more typical of human idiopathic ASD than those in monogenetic ASD rodent models ^22^. The characteristics of VPA-exposed marmosets suggest that these ASD models may be useful for bridging rodent ASD studies and human ASD.

In this study, we investigated the temporal remodeling of neural circuits using in vivo two-photon longitudinal imaging in VPA-exposed adult marmosets. We analyzed synaptic dynamics at 3-day intervals in the apical tuft dendrites of pyramidal neurons in the primate-specific dorsomedial PFC (dmPFC). The dmPFC is involved in social cognition and habit formation, and was found to exhibit less activity in nonverbal information-biased judgment in ASD individuals ^23^. We also observed the axonal boutons of local and long-distance cortical callosal connections labeled with fluorescent proteins of different colors to investigate whether circuit-specific synaptic remodeling was altered in the ASD model animal. Our study revealed that in VPA-exposed marmosets, turnover of postsynaptic dendritic spines was upregulated, and spines were actively generated in clusters but seemed to be randomly eliminated. Notably, in the marmoset model, the rates of both generated clustered spines and subsequently surviving spines (carryover spines) were much higher than in controls (3.3 and 5.9 fold, respectively). Presynaptic boutons of local axons, but not long-range commissural axons, showed more frequent turnover in model marmosets than in controls. Importantly, the clustering propensity of emergent spines in VPA-exposed marmosets was reduced by intranasal administration of oxytocin, which is currently being tested as a therapy for human ASD. These results suggest that both enhanced generation of clustered spines and altered turnover of local neuronal circuits are associated with ASD. We consider that these processes may be potential therapeutic targets for ASD symptoms.

## Results

### Two-photon imaging of the dendrites of PFC layer-2/3 pyramidal neurons in living marmosets

Changes in spine geometry (density) have been observed in the brains of individuals with ASD ^24^, and alterations in synaptic electrophysiological characteristics have been reported in animal models of ASD ^22, 25^. In the present study, we used a marmoset ASD model to investigate spine dynamics in the dmPFC. We used three adult VPA-exposed marmosets and four adult unexposed (UE) marmosets (See **Methods**). We inoculated the marmoset dmPFC with adeno-associated virus (AAV) and expressed fluorescent proteins mainly in layer-2/3 pyramidal neurons ^26^. These proteins showed sufficient fluorescence for in vivo two-photon imaging. The fluorescent protein-expressing neurons were locally distributed with an average diameter of 1.81 mm (range: 1.02-2.87) axially and 1.53 mm (range: 0.86-2.21) laterally. Post-experiment immunohistochemistry using antibodies against Iba-1 and GFAP showed no obvious increases in activated microglia or astrocytes, respectively (Figure S1).

We performed time-lapse observation of the dmPFC using a two-photon microscope ^26, 27^ (Figures 1A and 1B). Every 3 days, we conducted three imaging sessions of identical apical tuft dendrites in UE and VPA-exposed animals (*n* = 14 and 12 dendrites, respectively; mean dendrite span per branchlet: 31.7 ± 5.9 µm and 31.6 ± 5.8 µm, respectively). We found no significant differences in the dendritic spine density on apical tuft dendrites between the two types of marmosets (Figure 1C). We next compared the spine generation and elimination rates between the VPA-exposed and UE groups (Figures 1B and 1D). We found that the rates of both spine generation and elimination were approximately two times higher in VPA-exposed than UE marmosets (Figure 1D). This result was consistent with the results from various ASD mouse models of enhanced spine generation and elimination ^7–10^.

**Figure 1.**
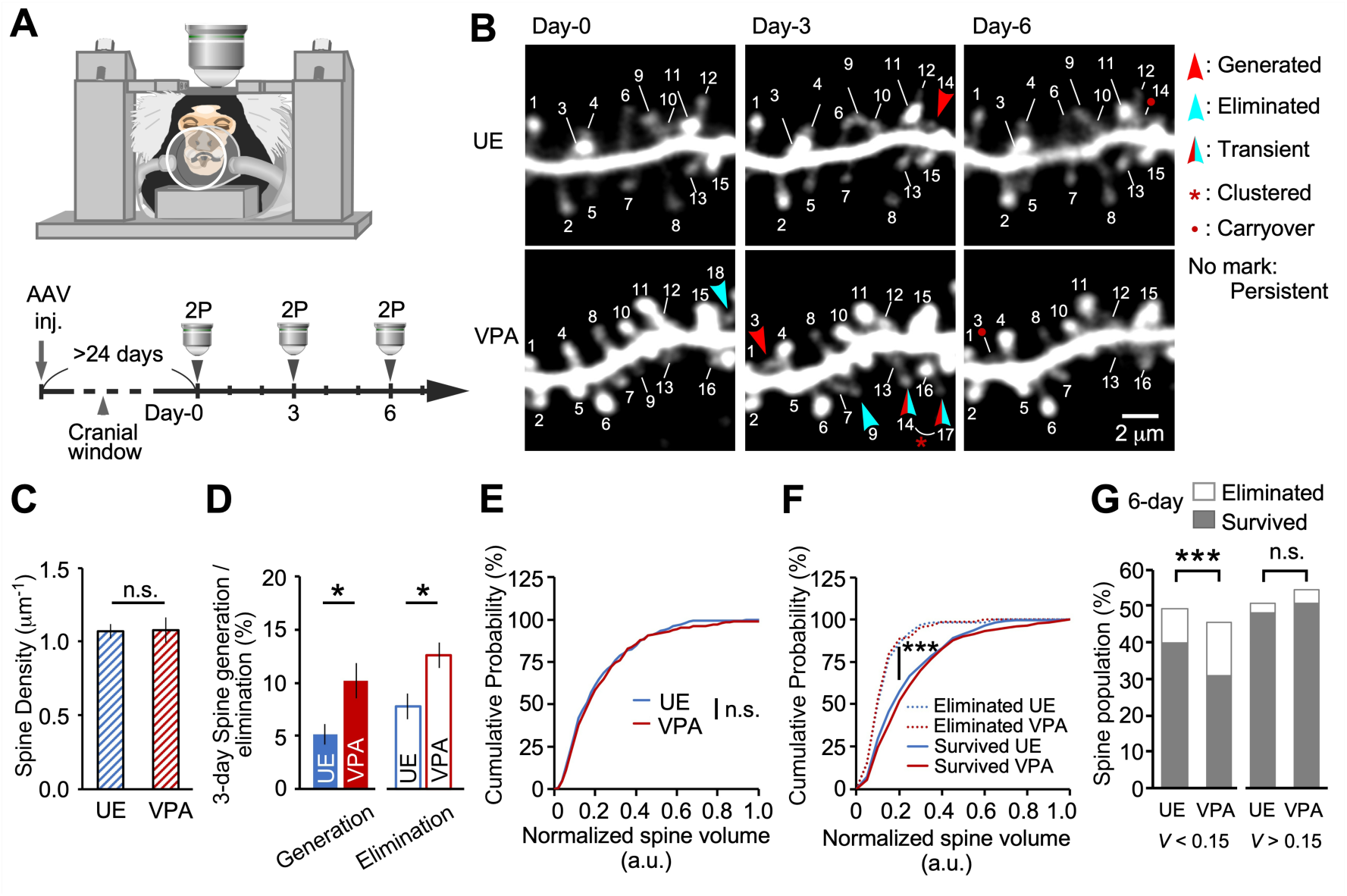
In vivo two-photon imaging of mature marmoset dendrites. **(A)** A marmoset was anesthetized for two-photon (2P) imaging. The lower panel shows the experimental schema. (B) Longitudinal 2P imaging from the PFC layer-2/3 pyramidal neuron tuft dendrites. Every 3 days, the same dendrites were imaged in UE (unexposed; control animals) and VPA-exposed (ASD model) marmosets, and the same spines were labeled using the same numbers throughout the period. (C) Mean spine densities of dendrites (mean ± s.e.m.; *P* = 0.85, Mann-Whitney U test; *n* = 14 and 12 dendrites in four and three UE and VPA-exposed animals, respectively). (D) Three-day spine generation rates (mean ± s.e.m.; *P* = 0.047, Mann-Whitney U test; the same dendrites as in (C)), and 3-day elimination *(P* = 0.044). **(E** and F) Cumulative plots of the normalized spine volume distribution (E) *(P* = 0.72; Kolmogorov-Smirnov test; *n =* 536 and 473 spines in UE and VPA-exposed animals, respectively), and cumulative plots of eliminated UE, eliminated VPA-exposed, surviving UE, and surviving VPA-exposed spines during the 6-day observation (F) *(P <* 0.001; *n* = 65, 87, 471, and 386 spines, respectively). (G) The population of eliminated and surviving spines during the 6-day observation are shown separately for groups with a spine volume of *V<* 0.15 and *V>* 0.15 *(V* <0.15; *P <* 0.001; Fisher’s exact test; *n* = 50, 69, 214, and 146 spines as in (F)) (F> 0.15; P = 0.59; *n* = 15, 18, 257, and 240 spines as in (F)). ***P < 0.001; *P < 0.05; n.s., not significant.

### Smaller spines in the ASD model marmoset were more prone to elimination

We next performed an analysis of spine volume, which is a crucial measure of synaptic weight. We calculated the normalized spine volume by dividing each spine’s fluorescent intensity with the dendrite shaft intensity, and pooled the results of all dendrites. The results showed no significant difference in the distribution of spine volume between the VPA-exposed and UE groups (Figure 1E). As in previous studies ^28, 29^, both groups showed a significant difference in volume distribution between spines that were eliminated during the 6-day period and those that survived (Figure 1F). The volume distribution of the VPA-exposed and UE groups did not differ when the eliminated and surviving groups were analyzed separately (Figure 1F); however, for the smaller spines, the proportion of eliminated spines was significantly larger in the VPA-exposed group (Figure 1G). These results suggest that the synaptic weight distributions in the dendrites were similar in the two types of marmosets, whereas the smaller spines were more vulnerable in VPA-exposed animals.

### Newly generated spines clustered more frequently in VPA-exposed marmosets than in controls

The generation of spines in close proximity to each other, or clustered spine generation, is considered to have functional significance, especially in learning and memory ^5, 10, 15, 18^. We next examined whether the generated spines in the VPA-exposed and UE marmosets were clustered or not (Figure 2A). We chose a 3 µm window for our analyses, since several biochemical, physiological, and structural studies have suggested that a 3 to 10 µm distance between spines facilitates sharing of resources, spine co-activation, and learning-induced structural plasticity ^30–36^. Moreover, the difference between the UE and VPA-exposed animals in spine clustering reached a plateau at a distance of 3 µm between spines, suggesting that events occurring within 3 µm are particularly important (Figure S2A). Clustered spine generation occurred 3.3 times more frequently in VPA-exposed animals than in UE animals (Figure 2B), even though the total number of generated spines in VPA-exposed marmosets was only about twice as many as that in the UE marmosets (Figure 1D). The clustered generated spines even accounted for 6.2% of the total spines on the dendrites. Next, we created a cumulative plot of the distance between generated spines. As shown in Figure 2C, there was a significant difference in the distribution of interspine distances between the VPA-exposed and UE groups. Again, the difference between the VPA-exposed and UE animals in terms of the probability of clustered spine generation was large (2.6 fold; Figure 2C). These results suggest the existence of a mechanism by which dendritic spines actively appear in clusters in the animals exposed to VPA. Therefore, we conducted a Monte Carlo simulation experiment to determine whether there was more clustering bias in VPA-exposed marmosets than in the hypothetical uniform random spine distribution (Figure 2D). To prevent underestimation of the inter-spine distances between the newly generated and eliminated spines, we conducted a simulation with all measured dendrites concatenated into one long dendrite, where dendrite lengths and spine counts of each dendrite were summed. The difference in clustering probability of the generated spine pairs between the UE and VPA-exposed animals was much larger than that of the eliminated spine pairs. In addition, the clustering probability of the generated spine pairs was greater than the 95^th^ percentile of the simulation distribution in the VPA-exposed group (Figure 2F), but remained below the 95^th^ percentile in the UE group (Figure 2E). The clustering probability of the eliminated spine pairs did not exceed the 95^th^ percentile in either the UE group (Figure 2G) or the VPA-exposed group (Figure 2H). Considering that the generated clustered spines may appear on specific dendrites, we conducted another simulation in which the dendrite length and the number of spines generated on each dendrite were left unaltered (that is, each dendrite was treated separately without concatenation), and obtained similar results (Figures S2D-H). We conclude that the clustering of newly generated spines, but not that of eliminated spines, on the dendrites of VPA-exposed marmosets is substantially enhanced rather than randomly distributed.

**Figure 2.**
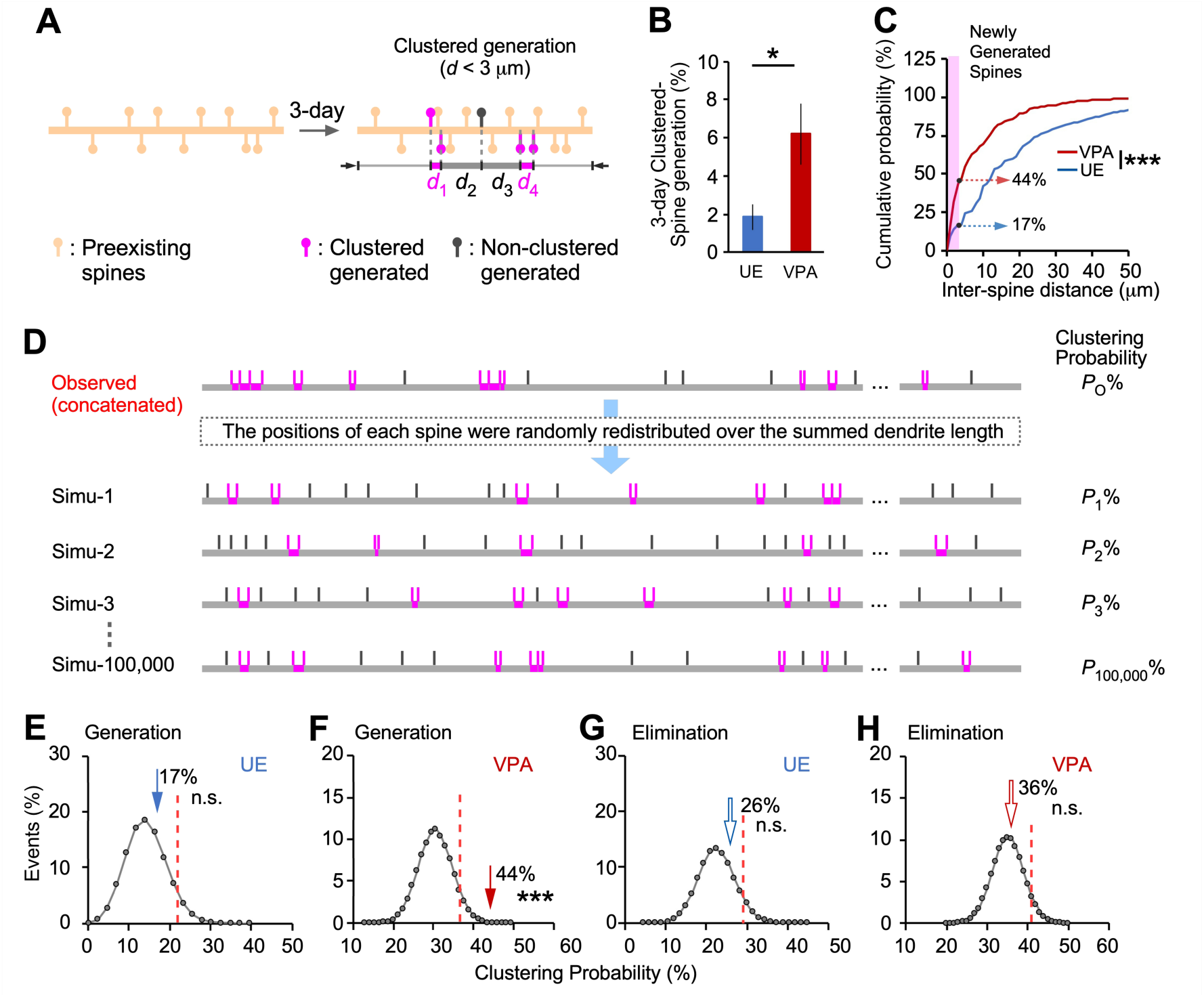
Dendritic spine clustered generation is more predominant than clustered elimination in VPA-exposed marmosets. **(A)** Schematic drawing of clustered spine generation. Pairs of newly generated or eliminated spines were considered to be clustered if they occurred within 3 pm of each other. Spines colored magenta or gray represent clustered and non-clustered spines, respectively. (B) Comparison of the percentage of clustered generated spines to the total number of spines between UE and VPA-exposed animals (mean ± s.e.m.; *P* = 0.016, Mann-Whitney U test; *n* = 14 and 12 dendrites in four UE and three VPA-exposed animals, respectively). (C) Distribution of inter-spine distances between newly generated spines *(P* = 0.0009; Kolmogorov-Smirnov test; *n =* 45 and 85 spine pairs in UE and VPA-exposed animals, respectively). Magenta-shaded area indicates inter-spine distances shorter than 3 pm, and numbers indicate the probabilities of clustering (within 3 pm). To prevent underestimation of the inter-spine distance, dendrites were concatenated into one long dendrite. (D) Validation of clustering bias by Monte Carlo simulation. In the simulation, the new spine positions were randomly determined with a uniform distribution over the summed dendrite length. Clustering probabilities for all inter-spine distances were calculated for each simulation, and the distributions of the clustering probability from 100,000 iterations are shown in **(E)-(H).** (E-H) Circles connected with gray lines represent probability plots of clustering events from 100,000 simulations; the actual numbers of spine clusters are represented by arrows (P = 0.23 and 0.00042; *n* = 44 and 86 newly generated spines in 14 and 12 dendrites in UE and VPA-exposed animals, respectively) (P = 0.20 and 0.36; *n* = 68 and 100 newly eliminated spines in UE and VPA-exposed animals, respectively). Dotted red lines show 95th percentiles. ***P < 0.001; *P < 0.05; n.s., not significant.

### The carryover fraction of newly generated clustered spines was higher in VPA-exposed marmosets than in controls

Three consecutive imaging sessions allowed us to monitor the fate of pre-existing and newly generated spines over 3 days ^9^. The carryover fraction was defined as the fraction of newly generated spines that survived to the last session (Figures 3A and 3B, GS). The carryover fraction was two times higher in VPA-exposed marmosets than in UE marmosets (Figure 3C). This suggests that experience-induced changes in spines may be more likely to persist in VPA-exposed marmosets. We further divided the newly generated spines into clustered and non-clustered spines and computed their carryover fractions. The carryover fraction of clustered spines in VPA-exposed marmosets was 5.9 times higher than that in UE marmosets (Figure 3D). On the other hand, there was no difference in the carryover fraction of non-clustered spines with or without VPA treatment (Figure 3D). Thus, clustered spines accounted for a greater proportion of spine carryover in the model marmosets than in the UE marmosets.

**Figure 3.**
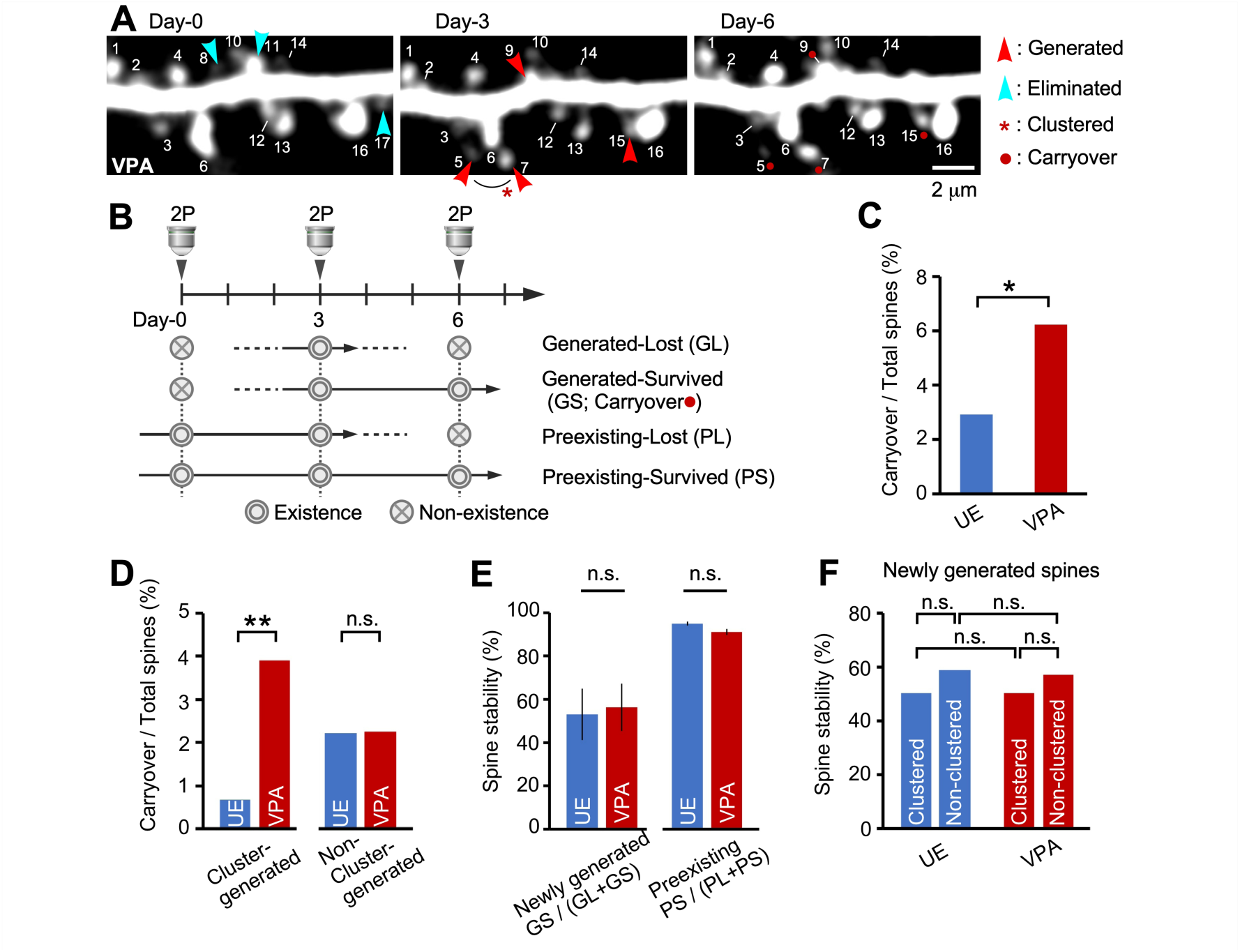
The surviving fraction of newly generated spines (carryover spines) is much larger in VPA-exposed marmosets than in controls, although spine stability is similar. **(A)** Representative dendrite images taken every 3 days are presented as in Fig. IB. **(B)** A diagram showing four patterns of spine generation and elimination. The red dot in the legend indicates the carryover spines labeled with a red dot in Figs. IB and 3A. (C) The ratio of the surviving fraction of newly formed spines (carryover spines) to the total spines was larger in VPA-exposed animals than in UE animals. *(P* = 0.037, Fisher’s exact test; *n* = 13, 448, 22, and 355 for UE carryover spines, UE total spines, VPA-exposed carryover spines, and VPA-exposed total spines, respectively). (D) Among clustered generated spines, the ratio of the carryover spine fraction to the total spines was much larger in VPA-exposed animals than in UE animals *(P* = 0.0022, Fisher’s exact test; *n* = 3, 448, 14, and 3 5 5 for UE carryover, UE total, VPA-exposed carryover, and VPA-exposed total spines, respectively). By contrast, the carryover spine fraction was not significantly different among non-clustered spines *(P* > 0.99, Fisher’s exact test; *n* = 10, 448, 8, and 355 for UE carryover, UE total, VPA-exposed carryover, and VPA-exposed total spines, respectively). **(E)** The stability of each newly generated and pre-existing spine was not significantly different between UE and VPA-exposed animals (mean ± s.e.m.; *P* = 0.94 and *P* = 0.091; Mann-Whitney U test; *n* = 13 and 10 dendrites from three UE and three VPA-exposed animals, respectively). (F) The stability of each newly formed spine was not significantly different between the clustered and non-clustered spines *(P >* 0.99; Fisher’s exact test; *n* = 3, 10, 3, and 7 for clustered GS, non-clustered GS, clustered GL, and non-clustered GL spines, respectively, in UE animals) *(P* > 0.99; Fisher’s exact test; *n* = 14, 8, 14, and 6, respectively, in VPA-exposed animals). *\*\*P* < 0.01; *\*P* < 0.05; n.s., not significant.

Having analyzed the collective nature of spines, we next analyzed the temporal stability of individual spines. Spine stability was defined as the percent ratio of newly generated or pre-existing spines that survived to the last session. Both in the VPA-exposed and UE groups, the spine stability of newly generated spines (∼50%) was considerably lower than that of pre-existing spines (∼90%) (Figure 3E). However, the stability of these two types of spines did not differ between the VPA-exposed and UE groups (Figure 3E). Furthermore, spine stability in both the VPA-exposed and UE groups was independent of the presence or absence of clustering (Figure 3F). The fact that spine stability was the same regardless of clustering suggests that interactions among newly formed spines have little effect on spine stability for some time after spine generation. In summary, clustered carryover spines comprised a much higher proportion of total spines in the VPA-exposed marmosets than in the UE animals, while the stability of individual spines was equivalent in the two groups.

Lastly, we compared the clustering of VPA-exposed marmosets with those of a mouse model. Exaggerated clustering of generated spines has been reported in a mouse model of *MeCP2* gene duplication, which results in a rare syndrome with core symptoms of ASD ^10^. The mean values of clustered emergent spines in the mouse study were obtained from the graphs in the paper. The results showed that the frequency of clustered emergent spines in the VPA-exposed marmosets was 9.4 per 100 µm, which was 3.9 times higher than that in *MeCP2* duplication mice (2.4 per 100 µm) (Figure S2B). Like the clustered spine generation rate, the carryover fraction in VPA-exposed marmosets was much higher than reported in *MeCP2* duplication mice. The frequency of carryover spines that appeared in clusters was 5.3 per 100 µm in the VPA-exposed marmosets, which was 5.7 times higher than in the *MeCP2* duplication mice (0.9 per 100 µm). Thus, the rate of spine clustering in the marmoset model differed markedly in comparison to the *MeCP2* duplication mouse model.

### Axonal boutons had a higher turnover rate in VPA-exposed marmosets than in controls

Deficits in projection-specific connectivity or cortical interaction have often been discussed in ASD, as exemplified by local overconnectivity and long-range underconnectivity ^37–39^. Spine turnover may depend on which neuron the coupled axons originate from ^7^. We next analyzed marmoset axons that were transfected with colored fluorescent proteins that differed according to the dmPFC hemisphere from which they projected (Figure 4A). We expressed the red fluorescent protein tdTomato in neurons ipsilateral to the observation window in the PFC, and the green fluorescent protein mClover in neurons on the contralateral side (same axial position and same distance from the midline; Figure 4A). Therefore, we were able to observe red dendrites and red axons from local neurons, as well as green axons from contralateral neurons (Figure 4B). In the magnified images, varicosities called presynaptic boutons were seen on the axons; new generation and elimination of these varicosities were seen on each of the three observation days (Figure 4C). Previous studies have shown that these boutons constitute presynapses ^40^, although some presynaptic terminals lack distinct varicosities (i.e., boutons) on the axons. In the present sample, most boutons were *en passant* boutons, and relatively few terminal boutons which had a neck between a varicosity and a parental axon shaft (Figure 4C). In this study, we considered the boutons to represent the existence of presynapses. We first measured the axonal bouton density and found a significant difference between the mean values of the ipsilateral and contralateral UE marmoset axons. By contrast, the difference between the ipsilateral and contralateral axons in VPA-exposed marmosets was not significant (Figure 4D). Next, we calculated the bouton size, which represents the synaptic weight ^40^, under the assumption that it was proportional to the maximum fluorescence intensity. We found that the distribution of bouton size was almost identical between the VPA-exposed and UE groups (Figure 4E). The size distribution was consistent in both spines and boutons, indicating that the balance of excitatory neuronal inputs is maintained in this autism model. We then calculated the 3-day bouton generation and elimination rates. Consistent with the rates of spine generation and elimination, those of bouton generation and elimination were higher in VPA-exposed animals (Figure 4F). Detailed analysis of each type of axon showed a significant difference in ipsilateral bouton gain between the two groups, whereas there was no significant difference in contralateral bouton gain (Figure 4G). This suggested that the difference in the generation and elimination of synapses between the VPA-exposed and UE groups may have been caused by the alteration of ipsilateral (i.e., local) axons. The 3-day bouton turnover rate was higher than that of spines (spine: 10.2% and 12.6% for VPA-exposed group, 5.2% and 7.8% for UE group, for generation and elimination, respectively). These results suggest that a mechanism that regulates appropriate synaptic remodeling in a projection-specific manner exists in control animals and that it is disrupted (hyperdynamic turnover in the local circuits) in the ASD model.

**Figure 4.**
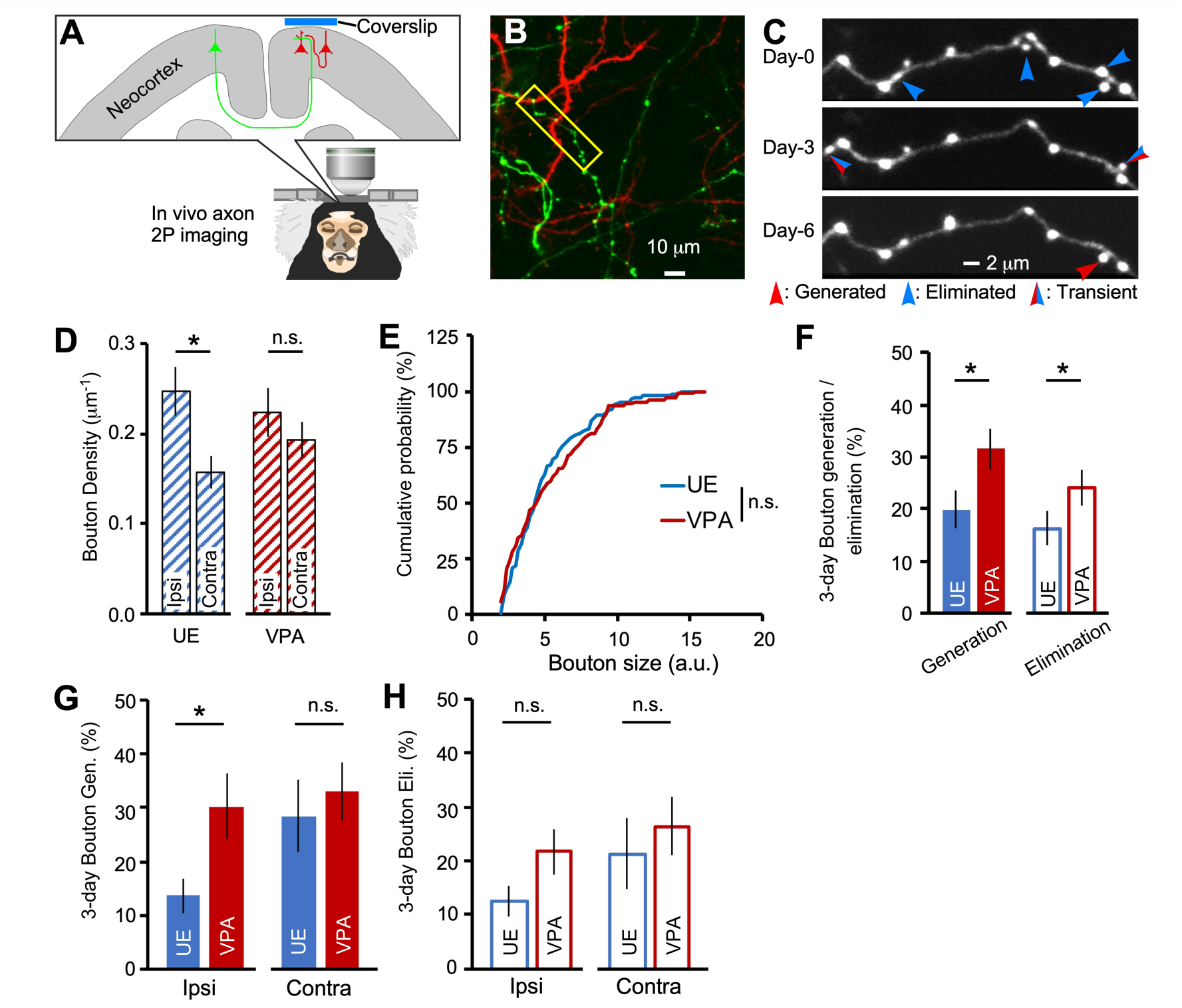
Two-photon in vivo axon imaging in ASD model marmosets. **(A)** AAV vectors expressing different colored fluorescent proteins were inoculated into each hemisphere of the dmPFC. (B) Axons from the contralateral hemisphere had green fluorescence, and axons from the ipsilateral hemisphere and dendrites had red fluorescence. **(C)** A representative axon from a VPA-exposed animal (the axon surrounded by the yellow rectangle in (B)) was imaged every 3 days. (D) Mean bouton densities are significantly larger in ipsilateral axons than in contralateral axons in UE animals *(P* = 0.035, Mann-Whitney U test; *n* = 28 and 21 ipsilateral and contralateral axons, respectively, from three animals). They are not significantly different between ipsilateral and contralateral axons in VPA-exposed animals (P = 0.36; h = 22 and 25 ipsilateral and contralateral axons, respectively, from two animals). The mean densities of terminal boutons were much lower than those of en passant boutons in our sample (mean terminal bouton density: 0.023, 0.012, 0.021, and 0.010 boutons/pm for UE ipsilateral, UE contralateral, VPA-exposed ipsilateral, and VPA-exposed contralateral axons, respectively). (E) The bouton size distribution between these animals (P = 0.57; Kolmogorov-Smirnov test; *n* = 149 and 107 boutons in 49 and 47 axons in three and two UE and VPA-exposed animals, respectively). (F) There is a significant difference in three-day bouton generation rates between these animals (P = 0.012, Mann-Whitney U test; *n =* 49 and 47 axons in three and two UE and VPA-exposed animals, respectively), and also in 3-day bouton elimination (P = 0.043). (G and H) Three-day mean bouton generation rates (G) (mean ± s.e.m.; *P* = 0.018 and *P* = 0.49, Mann-Whitney U test; *n* = 28, 22, 21, and 25 for UE ipsilateral, VPA-exposed ipsilateral, UE contralateral, and VPA-exposed contralateral axons, respectively) and elimination rates (H) (P = 0.093 and *P* = 0.28) for each axon type. *P < 0.05; n.s., not significant.

### Oxytocin nasal administration modified dendritic spine clustering

Oxytocin, a prosocial neuropeptide, is a potentially effective treatment for ASD symptoms. Oxytocin modifies the social skills of typically developed human individuals, and improves sociality in mouse ASD models ^41–43^; therefore, its potential as a therapeutic agent for ASD has long been anticipated. However, the benefits of oxytocin in human clinical trials are currently inconclusive ^44–47^. In order to better understand the clinical applications of oxytocin, it is therefore important to investigate its mechanism of action on synaptic phenotypes, which were characterized in this study in VPA-exposed marmosets. Thus, we next examined how the aforementioned properties of cortical spines changed after the administration of oxytocin. Oxytocin receptors have been shown to be expressed in the mouse cerebral cortex presynapse and postsynapse of putative excitatory neurons, but are not yet known in marmosets^48–50^. We have demonstrated the presence of the oxytocin receptor protein through Western blot analysis within the synaptosomal fraction isolated from the gray matter of the adult marmoset prefrontal cortex (Figure S5). Marmosets were administered saline and then oxytocin, both intranasally (Figure 5A). The average results from all marmosets failed to show a significant effect of oxytocin on spine generation and elimination rates (Figures 5B and S3A). Alternatively, analysis of the distance between generated spines suggested that oxytocin may affect the clustering efficiency of newly generated spines (Figures S3C and S3D). Thus, we next examined the effect of oxytocin administration on spine clustering (Figures 5C-J). After the oxytocin treatment period began, the proportion of clustered generated spines gradually decreased while that of non-clustered generated spines increased, although the difference was not significant (Figure 5C). We therefore tested whether the clustering bias was altered by the administration of oxytocin using simulations similar to those used in Figures 2D-H. The clustering bias between generated spines observed in VPA-exposed animals during saline administration was reduced by nasal oxytocin administration (Figures 5D-F, J, and S4). By contrast, oxytocin had a less pronounced effect on the proximity between eliminated spines (Figures S3B, 5G-I, 5J, and S4). We next analyzed the effect of oxytocin on axons. The bouton density, generation, and elimination of each axon species were not significantly different before and after oxytocin administration (Figure S6).

**Figure 5.**
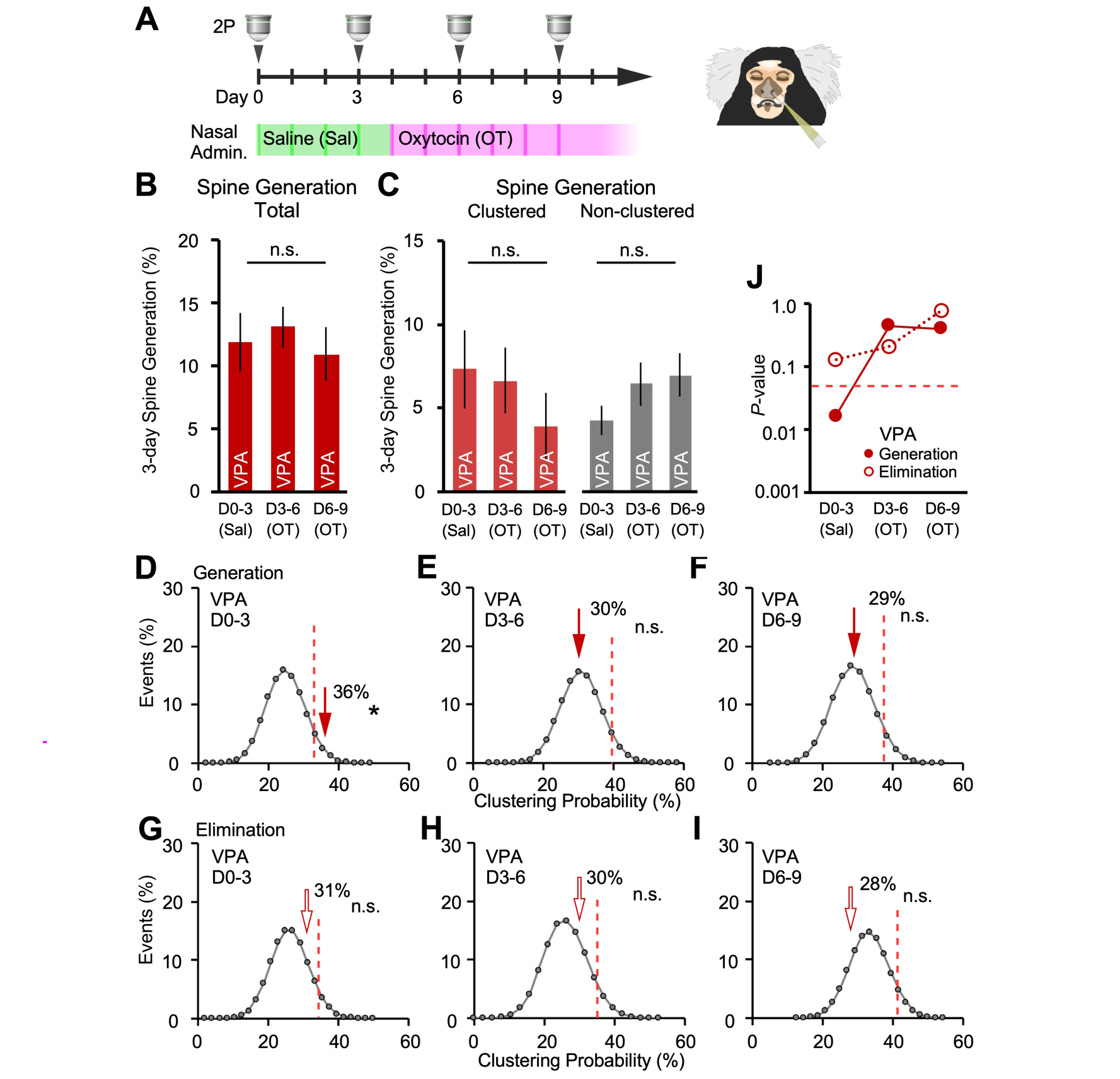
Oxytocin modifies the proximity of spine generation in model marmosets. **(A)** Schematics of oxytocin nasal administration and two-photon (2P) imaging (Left panel) and transnasal administration using a micropipette (Right panel). Saline was given to marmosets in the green-shaded period, while oxytocin was given in the magenta-shaded period. (B) Three-day spine generation rates were not modified after oxytocin administration (mean ± s.e.m.; *P* = 0.34, Friedman test; *n* = 12 dendrites in three VPA-exposed animals). **(C)** Three-day clustered or non-clustered spine generation rates were markedly (but not significantly) modified after oxytocin administration (mean ± s.e.m.; *P* = 0.22 and *P* = 0.56 for clustered and non-clustered spine generation, respectively, Friedman test; *n* = 12 dendrites in three VPA-exposed animals). **(D-I)** Effects of oxytocin on clustering bias of newly generated (D-F) and eliminated (G-I) spines. The graphs are shown as in Figs. 2E-H. *(n* = 45, 43, and 39 newly generated spine pairs and *n* = 48, 38, and 48 newly eliminated spine pairs during the DO-3, D3-6, and D6-9 periods, respectively, in three VPA-exposed animals). Dotted red lines show 95^th^ percentiles. (J) The P-values expressed in logarithm from (D)-(I) are indicated. *\*P* < 0.05; n.s., not significant.

### Gene expression analysis in adult ASD model marmosets

To explore the molecular mechanisms of altered spine dynamics in VPA-exposed marmosets and to assess the validity of adult VPA-exposed marmosets as a model of idiopathic ASD, we performed transcriptome analysis of the cerebral cortex using custom-made marmoset microarrays. Among the 9,296 genes expressed in cortical tissues, there were 2,484 differentially expressed genes (DEGs) with an adjusted *P*-value for multiple comparison (*P*_adj_) of < 0.05. First, we compared the DEGs in the cortical regions associated with social behavior between VPA-exposed marmosets and the postmortem brains of humans with ASD ^51^, and found a significant positive correlation between the two groups ^52^ (Figure 6A). This was similar to our previously reported similarity between juvenile VPA-exposed marmosets and humans with ASD ^22^. In addition, the modulation of gene expression in adult marmoset models and in human ASD was compared in each gene module that was previously configured by weighted gene co-expression network analysis (WGCNA) for samples from typically developed humans and those with ASD ^22, 52^. In both the marmoset model and human ASD, the majority of genes in modules associated with neurons and oligodendrocytes were downregulated (Figure 6B), while genes in modules associated with astrocytes and microglia were upregulated (Figure 6B). The direction of gene expression modulation of adult VPA-exposed marmosets was very similar to that of human ASD samples in 10 modules, compared to nine modules for juvenile VPA-exposed animals ^22^ (Figure 6C). The high similarity between VPA-exposed marmosets and human ASD contrasted with representative rodent monogenetic ASD models, which replicated human modulations only in limited gene modules and cell types (Figure 6C). One explanation for this may be that these transgenic mice are a model of a specific type of ASD in which one gene contributes particularly strongly, which is not the case in idiopathic ASD.

**Figure 6.**
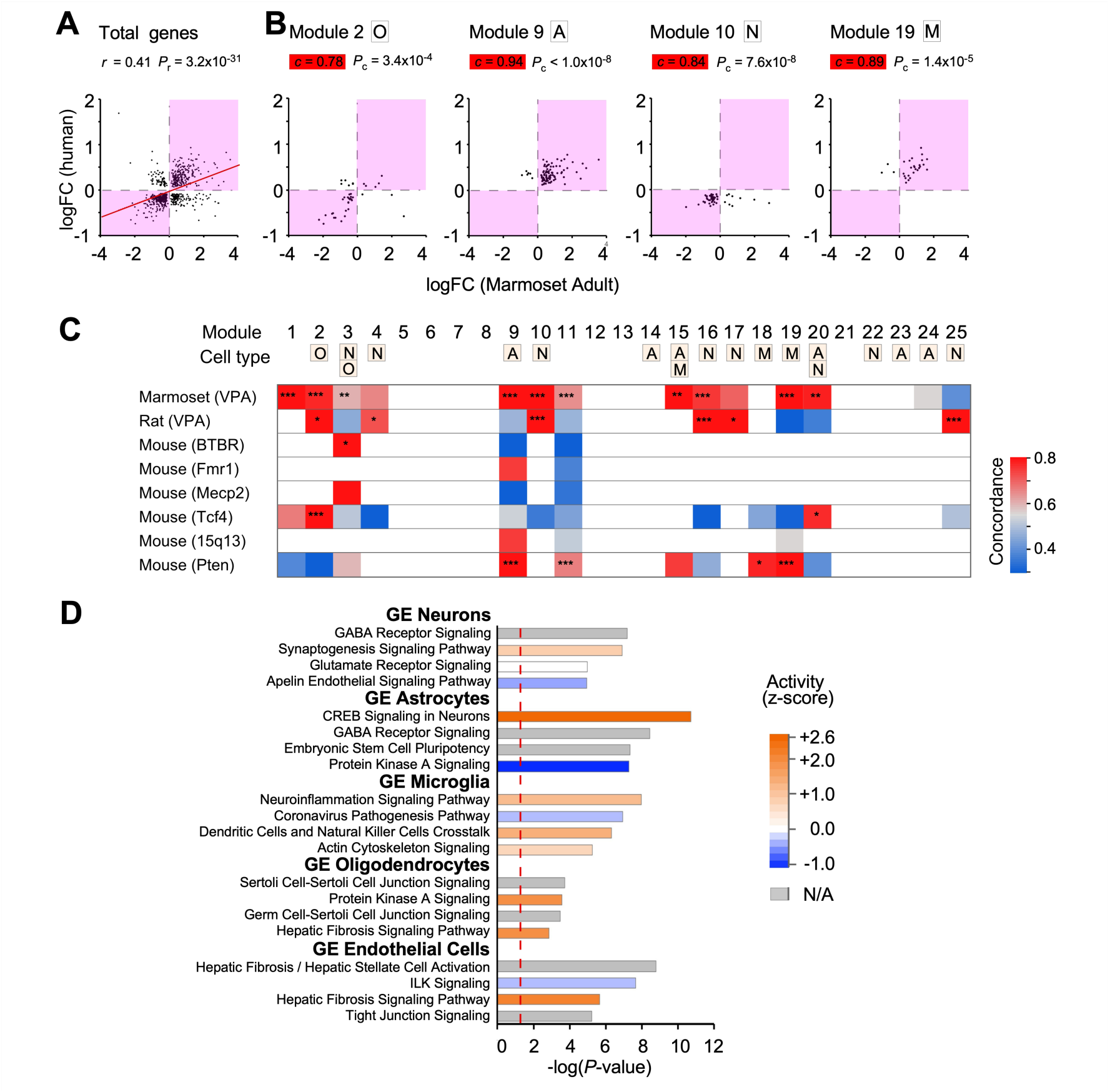
Gene expression modulations in adult marmoset social neocortical areas reflect those in human ASD as determined by microarray analysis. **(A, B)** Relationship of gene expression modulation between adult marmosets and postmortem samples from humans with ASD (A, B). As a measure of gene expression modulation, log fold-change (logFC) values were computed between ASD model animals and control animals, and then compared with human ASD data. Modulated genes with P <0.1 are plotted. Pearson’s correlation coefficient (r) and the P-values for the correlation coefficient *(P^)* are shown in (A). As a measure of concordance in the direction of gene expression modulation, one-sided P-values for concordance (P_c_) are shown in (B). The magenta-shaded areas represent the first and third quadrants, indicating that the two elements have changed in the same positive or negative direction. The red line in (A) is the linear regression. **(C)** Concordance of gene expression modulations across human modules. The color indicates concordance between animal model and human ASD modules with at least eight genes in common. Asterisks represent P-values determined using the one-sided binomial test: *P < 0.05, **P < 0.01, ***P < 0.001. For the rat VPA-exposed model and BTBR mice, genes with P <0.05 were selected; for other models and human ASD, those with P_ad_. < 0.1 were selected. (D) Enriched pathways for each cell type. The genes most closely associated with neurons, astrocytes, microglia, oligodendrocytes, and endothelial cells were analyzed. The color of the bars represents the direction of regulation of the pathway based on the logFC values. The P-values of enrichment were provided by the IPA software. The red dotted line represents the significance threshold (P = 0.05); N/A, not available.

We also analyzed DEGs in each cell type. This analysis confirmed the presence of a VPA-exposed marmoset-specific phenotype across the four cell types, as well as in vascular endothelial cells in the brain (Figure 6D). Pathway analysis of cell type-specific DEGs showed pathway abnormalities in each of these cell types. For example, microglial modules exhibited abnormalities associated with inflammation, which fits well with the inflammation hypothesis of ASD ^53^.

## Discussion

In this study, we used in vivo two-photon microscopy to monitor neurite dynamics in the apical tuft dendrites of dmPFC layer-2/3 pyramidal neurons and adjacent axons in VPA-exposed marmosets. The main findings were as follows. First, new spines were generated in clusters and subsequently carried over much more frequently than non-clustered spines. Second, turnover rates of both dendritic spines and presynaptic boutons were higher than in control animals. Specifically, bouton turnover in VPA-exposed marmosets was significantly greater in local axons, but not in callosal axons, compared to controls, indicating possible maladjustment of projection-specific circuit plasticity. Finally, nasal administration of oxytocin to model marmosets reduced the tendency of spines to cluster without affecting their turnover rate. These spine-associated effects of oxytocin support its therapeutic potential in individuals with ASD.

The process by which coactivating spines were formed in close proximity to each other during operant tasks was elucidated using a combination of behavioral tasks and functional two-photon imaging ^5^. Moreover, in layer-2/3 pyramidal neuron dendrites of the ferret visual cortex, neighboring spines tend to have similar visual representations ^54^. These and other reports suggest a link between the proximity of the distance between generated spines and similarities in their reactivity. In fact, electron microscopy studies have shown that adjacent spines often serve as postsynaptic targets for single axons ^55^. Clustered emergent spines that receive structurally or functionally similar inputs are thought to enable nonlinear dendritic computation and may facilitate the ability to acquire memory and behavioral skills through learning and experience ^11, 12, 14–17, 56^. Alternatively, clustered generated spines may also help stabilize formed circuits and accelerate learning due to their redundant involvement in the circuit ^57^. In fact, *CCR5* KO and *MeCP2* duplication mice commonly show elevated spine turnover and clustering as well as high learning efficiency ^10, 18^. On the other hand, over-efficient formation of neural circuits due to excessive synaptic clustering may result in the preservation of synapses that should be removed and may prevent the formation of additional neural circuits. Consistent with this hypothesis, VPA-exposed marmosets learned faster than UE marmosets during the first discrimination phase of a spatial reversal learning task, but learned slower after paradigm reversal ^58^.

Previous in vivo two-photon microscopy studies of ASD model mice have shown a common phenotype of high spine turnover in the motor and early sensory cortices that are less associated with ASD core symptoms than in the PFC ^7–10^. Accumulating evidence, including the results of this study, suggests that ASD-related genes and environmental factors converge across species, resulting in a phenotype of increased spine dynamics that continuously promotes neural circuit remodeling. Compared to spine turnover (generation and elimination), spine clustering in ASD animal models has been reported very infrequently to date, and only in the motor and visual cortices in *MeCP2* duplication models ^10^. Compared to VPA-exposed marmosets, the degree of clustered emergence in *MeCP2* duplication mice appears to be substantially lower. The reasons for this discrepancy are currently unclear. However, it may be due to differences in spine density, plasticity molecules, or subcellular organelles between species, cortical areas, syndromic and idiopathic ASD models, and deep and upper-layer pyramidal neurons. In any case, the strong clustering tendency in the dmPFC of VPA-exposed marmosets underscores the importance of clustering in the pathogenesis of idiopathic ASD and warrants further research of clustered spine emergence in both physiological and pathophysiological conditions.

The synaptic connections of local neural circuits exhibited more plasticity in VPA-exposed animals than in UE animals, whereas the long-distance callosal neural circuits did not show plasticity differences. This suggests that projection-specific plasticity control may be altered in ASD ^37, 38, 59^. The excessive plasticity in the local neuronal circuits of VPA-exposed marmosets is reminiscent of the enhanced long-term potentiation of layer-5 pyramidal neurons in acute slice preparations in a VPA-exposed rat ASD model ^60^. However, differences in synaptic remodeling of different types of cortical inputs have never been compared in living animals or in vitro in ASD models. Postsynaptic neuron-specific deletion of the *Fmr1* gene in mice reduced the amplitude of synaptic transmission in callosal inputs but not in local inputs, suggesting the existence of differential regulation between the group of inputs ^61^. Several experiments, including single-cell transcriptome analyses, have shown a variable number of DEGs between cortical layer-specific excitatory neurons^62^ or between cortical areas ^51, 63^ in human ASD. These differences may serve as the basis for projection-specific circuit plasticity. Abnormalities in brain connectivity strength between ASD-relevant brain regions in humans with ASD have been demonstrated by functional brain imaging techniques such as functional magnetic resonance imaging (fMRI) and magnetoencephalography (MEG) /electroencephalography (EEG), using the degree of brain activity coherence as an indicator ^1, 38, 40^. Differences in the degree of bouton dynamics between local and callosal connections in VPA-exposed marmosets suggest that in addition to variations in connection strength, changes in projection-specific synaptic remodeling may also contribute to ASD symptoms. It will be interesting to analyze whether mismatches in plasticity control involving local and long-range connections facilitate autonomous local circuit remodeling independent of the highly contextualized information computed in the brain-wide global network.

To identify therapeutic targets, it will be valuable to determine the molecular mechanisms that govern spine turnover and spine clustering in VPA-exposed marmosets ^64^. Our study demonstrated that oxytocin reduced clustering of generated spines in ASD model marmosets without altering spine turnover, while in mice the effect of oxytocin on synaptic plasticity has been studied in various brain regions such as the olfactory system, hypothalamus, and neocortex ^42, 65^. In the present study, we have substantiated the presence of the oxytocin receptor protein through Western blot analysis within the synaptosomal fraction isolated from the gray matter of the adult marmoset prefrontal cortex (Figure S5). This observation implies that chronic oxytocin exposure potentially exerts a direct inhibitory influence on spine clustering by modulating intracellular signaling pathways, notably the MAPK/ERK pathway, downstream of oxytocin receptor activation. Oxytocin receptors in marmosets are notably expressed in the basal ganglia of the nucleus basalis of Meynert (NBM), a prominent cholinergic region within the brain. The NBM serves as a significant source of cholinergic innervation to various brain regions, including the neocortex, and plays a pivotal role in regulating selective attention and motivational processes^50^. It is plausible that the alteration of spine clustering induced by oxytocin may involve the modulation of cholinergic inputs. Oxytocin’s inhibitory effect on spine clustering could restrain excessively prolonged circuit stability and mitigate behavioral perseverance, as reported in human ASD ^66^. Differences in oxytocin systems between rodents and primates may significantly affect therapeutic efficiency, which underscores the importance of primates in translational research on oxytocin ^50^. Spine clustering in the *MeCP2* duplication mouse model was suppressed by an inhibitor of Ras-MAPK(ERK) signaling without altering the spine generation rate ^67^. Ras, like other proteins such as Rho, Rac, and cofilin, has been reported to diffuse from activated spines to neighboring spines via dendrites and is thought to alter the plasticity of neighboring spines ^5, 31, 32, 68^. Interestingly, transcriptome analysis of our model marmosets showed four times greater upregulation of *TGF-β2*, which is upstream of the Ras-MAPK pathway, than control marmosets. Synaptic plasticity involves not only neurons but also diverse glial cells. We found that microglia demonstrated activation of signals related to viral infection, inflammation, and reorganization of the actin cytoskeleton (Figure 6D). Suppression of excessive microglial activation with minocycline or other matrix metalloproteinase-9 inhibitors has been reported to suppress excessive spine turnover in fragile X model mice ^8^. Our transcriptomic analysis also revealed significant downregulation of myelin-related genes, such as PLP1, which suppresses heightened cortical plasticity. On the other hand, it has been shown that the stress hormone corticosterone increases spine turnover in mice ^69^. Indeed, as in mouse ASD strains ^70^ and humans with ASD ^71^, our ASD model marmosets have aberrant cortisol responses ^72^, suggesting that environmental factors such as anxiety and endocrine system disorders may be responsible for the increase in synaptic remodeling. Therefore, oxytocin, which reduces spine clustering, could be beneficial for individuals with ASD when combined with the suppression of spine turnover.

### Limitations of the study

In this study we did not investigate inhibitory synapses. The location of inhibitory inputs and the local balance between excitatory and inhibitory inputs may also have a significant effect on nonlinear information integration by dendrites. We were also unable to analyze the genetic, anatomical, or electrophysiological characteristics of the pyramidal neurons that we examined. Activity-dependent labeling of neurons would allow for better characterization of the relationship between neuronal activity and spine clustering. Electron microscopy analysis of clustered generated spines would have made it possible to confirm the presence of common presynaptic partners. We believe that the future implementation of these approaches will better clarify the nature of spine clustering in autism models.

## Acknowledgments

We thank H. Kasai of The University of Tokyo for helpful discussion and suggestions about two-photon imaging, and both H. Yamasue of Hamamatsu University School of Medicine and T. Minamimoto of QST Japan for their careful reading and thoughtful comments on the manuscript. We are also grateful to T. Araki, S. Wakatsuki and M. Iwasaki of the National Center of Neurology and Psychiatry (NCNP) for helpful discussion about experiments and marmoset surgery. We also thank A. Tsuchiya, M. Nakamura, T. Sato, W. Suzuki, A. Mishima, K. Mimura, and N. Miyakawa of our lab for supporting primate experiments, and R. Saito, Y. Katakai, and other members of the NCNP primate facility for caring for VPA-exposed and UE marmosets. A. Sawatari and S. Ikeda of Iwate University assisted with spine volume analysis. English editing was performed by Zenis, co. Ltd., Kyoto, Japan. This work was supported by Intramural Research Grants for Neurological and Psychiatric Disorders from the NCNP (3-5, J.N.; 29-6, N.I.), a Novartis Research Grant 2019 (S.W.), JSPS KAKENHI Grant Numbers JP18K06497 and JP22K07363 (J.N.), and AMED Grant Number JP21dm0207066 (N.I.).

## Author contributions

J.N. and N.I. designed the study; J.N., S.W., R.I., and K.S. performed the in vivo two-photon imaging; K.N. generated the autism model marmosets; J.N. and R.I. conducted image analyses; E.S., H.T., H.M., A.W., and T.Y. prepared the AAV vectors.; S.W., T.O., K.S., K.H., K.S., and I.M. conducted the microarray analysis.; K.S. and R.I. performed immunohistochemistry; and J.N. and N.I. wrote the manuscript. All authors approved the final version of the manuscript. Correspondence and requests for data should be addressed to J.N. or N.I.

## Declaration of interests

The authors declare no competing interests.

## Inclusion and diversity

We support inclusive, diverse, and equitable research conduct.

## METHODS

### Animals

All experimental procedures were approved by the Animal Research Committee of the National Center of Neurology and Psychiatry. Common marmosets (*Callithrix jacchus*) were housed in captivity at the National Center of Neurology and Psychiatry under a 12-h/12-h light/dark cycle and were fed food (CMS-1; Clare Japan) and water ad libitum. Temperature was maintained at 27-30°C and humidity at 40-50%.

### Marmoset ASD model

Serum progesterone levels in female marmosets were regularly measured to determine the date of fertilization. Four percent VPA dissolved in 10% glucose solution was administered intragastrically to pregnant marmosets at 200 mg/kg/day daily for 7 days starting on gestation day 60. No doses were given to control mothers. The obtained ASD model offspring from VPA-administered mothers and control offspring from UE mothers were kept in their home cages until the day of inoculation with AAV vector. We used four UE marmosets (monkeys KR (f, 1.3), TR (f, 3.2), VR (m, 2.2), and BH (m, 1.3)) and three VPA-exposed marmosets (monkeys PR (f, 1.5), SH (f, 1.4), and MS (m, 1.3)); f or m and numbers in parentheses indicate sex and age in years at the time of AAV inoculation, respectively.

### AAV vector inoculation of the marmoset neocortex

Each experimental marmoset was pretreated with the antibiotic cefovecin sodium (8 mg/kg body weight, intramuscular (i.m.); Zoetis), prednisolone (2 mg/kg body weight; i.m.), the analgesic ketoprofen (2 mg/kg body weight, i.m.), and atropine (0.15 mg/kg body weight, i.m.). It was then anesthetized with ketamine (15 mg/kg body weight, i.m.; Daiichi-Sankyo) and xylazine (1.2 mg/kg body weight, i.m.; Bayer), which were supplemented with sevoflurane inhalation (2-4%). The marmoset was then fixed to a stereotaxic instrument (SR-6C-HT; Narishige, Tokyo, Japan). During all surgical procedures, marmosets were also supplied with humidified oxygen as needed, and warmed to 37-39 °C with a heating pad (FST-HPS; Fine Science Tools, North Vancouver, Canada). Body temperature, SpO_2_, heart rate, cardiac electrogram, respiratory rate, and actual concentration of oxygen and sevoflurane were measured using a biomonitor (BSM-5132; Nihonkoden, Tokyo). After incision of the skin at the midline, the skull was exposed and a 1-mm-diameter hole was made bilaterally above the PFC Brodmann area 8 (11.5 mm anterior to the interaural line, 3 mm lateral to the midline) using a dental drill.

A puller (PC-100; Narishige) was used to create a micropipette (tip diameter 30-50 µm) from a micro glass tube (1.0-mm diameter; TW100F-4; WPI, USA). The pulled micropipette was then beveled using a micro grinder (EG-402; Narishige) and was sterilized by overnight exposure to UV light or ethylene oxide. The pipette was set to a glass microsyringe (Model 1701,10 µL; Hamilton, USA) using a Priming kit (55750-01; Hamilton), and the microsyringe was back-filled with silicon oil and attached to a microsyringe pump (LEGATO 130; WPI) fixed to the manipulator of a stereotaxic instrument (SM-15R; Narishige). The micropipette was then tip-filled with AAV vector suspension and slowly inserted through the dura and into the neocortex, avoiding large blood vessels. The total 0.5-µL volume of AAV vector was then inoculated at 0.1 µL/min at a depth of 0.8 mm from the surface. The viral preparations were adjusted to the following final concentrations: 1×10^9^ vg/mL for rAAV1-Thy1S-tTA and 1×10^12^ vg/mL for AAV1-TRE3-tdTomato, expressing red fluorescence protein; 5×10^9^ vg/mL for rAAV2/1-Thy1S-tTA and 1×10^12^ vg/mL for rAAV2/1-TRE3-mClover, expressing green fluorescence protein. After the 5-min incubation period, the micropipette was retracted, a silicon plug was stuffed into the cranial hole, and the skin incision was closed with sutures. After recovery from anesthesia, the marmoset was returned to its home cage.

### Imaging window preparation

Each experimental marmoset was pretreated with the antibiotic cefovecin sodium, prednisolone, the analgesic ketoprofen, and atropine. It was then anesthetized with ketamine and xylazine, which were supplemented with sevoflurane inhalation (2-4%). The marmoset was then fixed to a stereotaxic instrument. The marmoset was also supplied with humidified oxygen and warmed up to 37-39 °C. After skin incision at the midline, the skull was exposed and a 4-mm-diameter hole was made at the point where the AAV was inoculated using a dental drill fixed to a stereotaxic manipulator (SM-15R; Narishige). The dura mater was carefully removed using microsurgical ophthalmic scissors, fine forceps, and a microhook to minimize any pressure applied to the surface of the brain. A piece of coverslip, consisting of three laminated circular coverslips (5-mm diameter + 3-mm diameter × 2), was used to cover the exposed brain surface^26^. The coverslip was fixed using dental acrylic (Fuji-Lute; GC Corp., Tokyo, Japan) to create an imaging window in the skull. A custom-made stainless-steel recording chamber (ICM, Tsukuba, Japan) was then attached to the skull using dental resin (Bistite II; Tokuyama Dental, Japan) such that the imaging window was in the center of the chamber. After the imaging window was covered with an acrylic lid, the marmoset was allowed to recover from the anesthesia in the monkey ICU room and was then returned to its home cage.

### In vivo two-photon excitation microscopy

In vivo two-photon imaging of neurites was performed using an upright microscope (BX61WI; Olympus) equipped with a laser scanning microscope system (FV1000; Olympus) and a water-immersion objective lens (XLPlanN, 25×, NA 1.05; Olympus). The system included mode-locked, femtosecond-pulse Ti:sapphire lasers (MaiTai; Spectra Physics) set at a wavelength of 980 nm.

Imaging sessions were conducted every 3 days. A marmoset was pretreated with atropine and anesthetized with sevoflurane inhalation (3-5%). The marmoset was laid in the prone position and its head was fixed using the imaging chamber and brass posts (Figure 1A). In each case, the imaging location was identified based on blood vessel morphology. Tuft dendritic branches (< 100 µm depth) of layer 2/3 pyramidal neurons were used for two-photon imaging experiments. An objective lens correction collar was manually set just before imaging so as to minimize spherical aberrations (i.e., to acquire the brightest image possible). The reciprocal scan mode was used to scan each *xy*-image (256 by 256 pixels; 65 msec/frame). Three-dimensional fluorescent images with 51 *xy*-images, each separated by 0.5 µm, were obtained at each imaging site.

### Dendritic spine and axonal bouton analysis

We applied a 2-pixel spatial Gaussian filter and then registered the obtained *xyz*-images using the StackReg or Multi-StackReg plug-in function of Image-J (Fiji)^73^. Spines were identified as protrusions from dendrites with an apparent head structure. Filopodial protrusions (which have no apparent spine head) were excluded from the analysis. Two spines observed during different sessions were considered to represent the same spine if the initial position subsequently exhibited the same distances from adjacent landmarks. A spine was interpreted as lost if it temporarily disappeared. The minimum spine length was set at 0.4 µm. Snowman-shaped spines (shaped like two spheres stuck together) were interpreted as two separate spines. Only spines that appeared laterally were included in the analysis (this underestimated the spine density). Dendrites that were visible in all sessions were used in the analysis.

We also analyzed the axons that appeared in the same images as the dendrites. For axon analysis, we used five marmosets that expressed mClover in the right hemisphere and tdTomato in the left hemisphere (UE monkeys: TR, VR, BH; VPA-exposed monkeys: SH, MS). We applied a 2-pixel spatial Gaussian filter to the *xyz*-images and obtained *z*-substacked average images for a predetermined z number for each axon. By combining images obtained at the optimal depths, we created montage images for each axon, avoiding dendrites and other axons present at other imaging depths. We then examined the brightness along the axon shaft using the Plot-profile function of Image-J^73^. The ratio of bouton brightness to neighboring axon shaft brightness was then calculated. We did not see axons that were much thicker than the diffraction limit of our two-photon imaging, and discarded areas that overlapped with other dendrites and axons. Boutons were detected as bright swellings along axons, with intensity values at least two-fold higher than that of the flanking axon backbone ^74^. Since we could not detect axonal synapses below the bouton threshold, the subthreshold boutons may have been misidentified as newly generated boutons when they enlarged and therefore exceeded the threshold in the next imaging session. As a consequence, the number of newly generated boutons may have been overestimated. Likewise, the number of boutons that were eliminated may also have been overestimated. This may explain the result that the overall bouton turnover was higher than the spine turnover. The 3-day bouton generation and elimination for unit axon length were obtained by dividing the number of boutons generated and eliminated over a 3-day period by the axon length. The 3-day bouton generation and elimination as a percentage were calculated by dividing the 3-day bouton generation and elimination for unit axon length by the mean bouton density. We did not exclude varicosities of transfer vesicles. In the next session, boutons located within 1 µm of the expected point were interpreted as identical to those in the previous session.

The percentage of newly formed spines or boutons was calculated as the number present at time point 2 but not at time point 1, divided by the total number present at time point 1 and multiplied by 100. The percentage of eliminated spines or boutons was calculated as the number present at time point 1 but not at time point 2, divided by the total number present at time point 1 and multiplied by 100.

### Monte Carlo simulation of newly generated or eliminated dendritic spines

To obtain unbiased distributions of the inter-spine distance of newly generated or eliminated spines, each dendrite was concatenated in random order, producing one long dendrite repeatedly (1,000 times permutation; Figures 2C and S3C, D). The average clustering probability was then calculated. We set the clustering threshold at 3 µm since several biochemical, physiological, and structural studies have suggested that a 3 to 10 µm distance between spines facilitates sharing of resources, spine co-activation, and learning-induced structural plasticity ^30–36^. In Fig. S2A, the threshold applied for the analyses (3 µm) was confirmed to be appropriate because it is located at the lowest end of the plateau ^10^. Monte Carlo simulation was performed to determine whether a clustering bias existed in the process of spine generation or elimination ^18^. The simulation used the total dendrite length and the total number of generated and eliminated spines. The positions of generated and eliminated spines on the dendrites were then regenerated with a uniform random distribution using Excel software. The distances between generated and eliminated spines were computed for all dendrites and the percentage of clustered events was calculated (Figure 2D). As clustered spines may appear more on specific dendrites, another simulation was conducted on individual dendrites without connecting each dendrite, and similar results were obtained (Figure S2D-H). The above simulations were performed with 100,000 iterations to obtain the distribution of clustering events, and the results are shown in Figures 2E-H, 5D-I, S2E-H, and S4A-F by connected gray lines.

### Dendritic spine volume measurement

We calculated the spine-head volume by partially summing the fluorescence values of five sequential *z*-images by taking the moving average of the image stack along the *z*-plane. This was done to avoid summing other dendrites or axonal fibers present in different imaging planes. Because the thickness of dendritic spines is near the diffraction limit of a two-photon microscope, partially summed values (2 µm range in the *z*-direction) can be used to reflect spine volumes. Thus, the maximum value of *z*-moving average images allowed us to obtain good approximations of the total *z*-summed stacked images. The normalized spine volume was obtained by dividing the spine fluorescence intensity by the fluorescence intensity of the adjacent dendrite shaft.

### Nasal oxytocin administration to VPA-exposed marmosets

New World monkey oxytocin (Pro-8 oxytocin ^75^) was custom synthesized (GenScript, Piscataway, NJ, USA) and was dissolved in 0.5 mg/mL of 0.9% NaCl solution. The oxytocin solution was typically administered 2.5 h before the two-photon imaging and at approximately same time on days without two-photon imaging. Oxytocin solution was applied to the nasal cavities of each marmoset at a dose of 150 µg/kg body weight as in a previous study^75^, using a micropipette equipped with a soft silicon tube on the tip. Oxytocin was administered every day (monkey MS) or every other day (monkeys PR and SH) starting on day−4 (Figure 5A). Verification of oxytocin delivery to the cerebral interstitial fluid was verified by measuring the plasma Pro8-oxytocin concentration using a sandwich competitive chemiluminescent enzyme immunoassay (CLEIA) (ASKA Pharma Medical, Fujisawa, Japan). The Pro8-oxytocin concentration in all blood samples was monotonically increased 30 min after nasal oxytocin administration (mean ± s.d. = 30.6 ± 14.8 pg/mL; *n* = 2 males and 3 females) but not 30 min after nasal saline administration (mean ± s.d. = 9.9 ± 1.7 pg/mL; *n* = 2 males and 3 females); thus, we concluded that Pro8-oxytocin was successfully delivered to the interstitial fluid of the neocortex.

### Microarray gene expression analysis

Microarray analysis of marmoset gene expression was performed using brain samples from three adult VPA-exposed and three UE marmosets, as previously described^22^. To increase data reliability, we pooled the data from the social brain areas (area 12 and area TE) that are affected in humans with ASD. Tissue sampling and GeneChip analysis were conducted as previously reported^22^. Briefly, marmosets were anesthetized with ketamine hydrochloride (50 mg/kg, i.m.) and sodium pentobarbital (90-230 mg/kg, intraperitoneal; Kyoritsu Seiyaku). After reflections had completely disappeared, the animals were transcardially perfused with diethyl pyrocarbonate-treated phosphate-buffered saline, and the cortical tissue was isolated and immersed in RNAlater (Thermo Fisher Scientific). Total RNA was extracted using the RNeasy Mini Kit (Qiagen, Hilden, Germany). RNA integrity was assessed using a Bioanalyzer (Agilent Technologies), and samples with RNA integrity number values >7 were evaluated. Biotin-labeled cRNA probes were prepared using the GeneChip 3’IVT Express Kit (Affymetrix). The probes were hybridized to a custom-made microarray (Marmo2a520631F) using the GeneChip Hybridization, Wash, and Stain Kit (Affymetrix). Microarrays were scanned using a GeneChip Scanner 3000 (Affymetrix) and processed using MAS5, and the reliability of probe detection was examined. Data were normalized using GCRMA. Genes with log2 expression values greater than 5 were considered to be expressed in brain tissue. The log fold-change (logFC) value was used as a measure of gene expression modulation and evaluated using Welch’s t-test with Benjamini-Hochberg adjustment (*P*_adj_). For affected genes with multiple probes, the data from the probe with the lowest *P*_adj_ was used. To compare gene expression modulations in the marmoset model with those in human ASD postmortem samples ^51^ commonly modulated genes with *P*_adj_ < 0.1 were used.

### Comparison of log fold change (logFC) values between human ASD and animal models, as determined by gene co-expression modules

To compare gene expression changes between the ASD group and the typically developed human group, or between the ASD model animals and controls, logFC values were calculated for the genes analyzed by the gene chip. The concordance values were then calculated for each gene module ^51, 52^ to show consistency in the direction of gene expression changes between human ASD and the animal models (Figure 6C). The concordance value was the percentage of genes that changed in a common direction (first and third quadrants in Figure 6B) in human ASD and the model animal. For modules with at least eight common genes between the animal model and human ASD, the concordance values are shown in color in Figure 6C. The logFC values of VPA rats^76^ (age 35 days), BTBR mice ^77^ (age 4 months), *Fmr1* KO mice^78^ (age 8-14 weeks), *MeCP2* heterozygous mice ^79^ (age 5 weeks), *Tcf4*^tr/+^ mice ^80^ (age 60-80 days), 15q13.3 deletion mice ^81^ (age 10-22 weeks), and *Pten*^m3m4^ mice ^82^ (age 6 weeks) were obtained from previously published studies. To compare gene expression modulations in the marmoset model with those in postmortem samples of humans with ASD ^51^, commonly modulated genes with *P*_adj_ < 0.1 were used. For the rat VPA models and BTBR mice, genes with *P*_adj_ < 0.05 were selected; for other models and human ASD, genes with *P*_adj_ < 0.1 were selected. To compare gene expression modulations in the rodent models with those in human ASD, gene symbols were converted using HomoloGene (NCBI, https://www.ncbi.nlm.nih.gov/homologene, release 68). Due to the small number of genes that are commonly affected, mouse models of maternal immune activation and *Shank3* knockout were not included in the list.

### Signaling pathway estimation of marmoset genes

Pathway analysis of marmoset genes was conducted using IPA software (Qiagen, Summer Release 2020). Genes specific to brain cell types (neurons, astrocytes, microglia, oligodendrocytes, and endothelial cells) were selected from genes with mean logFC gene enrichment values > 2, among the top-ranked cell type-enriched genes based on the human single-cell analysis data in a previous report ^83^.

### Examination of glial cell activation

Marmoset brains fixed in 4% PFA were replaced with 30% sucrose in 0.1 M phosphate buffer (pH 7.4). Frozen sections with 50 µm thickness were prepared using a microtome (Retoratome REM-710; Yamato-Kohki, Saitama, Japan). Brain slices were treated in antigen-retrieval solution (HistoVT One; Nacalai Tesque, Kyoto, Japan) at 70°C for 20 min, followed by PBS washing and blocking (blocking buffer: 5% normal goat serum, 0.4% TritonX-100 in PBS) at room temperature for 60 min. After blocking, free-floating brain slices were incubated in a solution of anti-Iba1 antibody (1:2000; Fuji film-Wako) or anti-GFAP antibody (1:10,000; Agilent-Dako) at 4°C overnight. The slices were washed with PBS (10 min, three times) and incubated with biotin-labeled goat anti-rabbit secondary antibody (1:500; Vector Labs) at room temperature for 90 min. The slices were washed with PBS (10 min, three times), then reacted with the ABC complex for 90 min at room temperature (Vectastain Elite ABC kit; Vector Labs). After washing with phosphate buffer (10 min, three times), the slices were mixed with nickel-sensitized DAB reaction solution (0.01% 3,3’-diaminobenzidine-4HCl) to develop color. The samples were washed with phosphate buffer, mounted on glass slides, and dehydrated with ethanol and xylene. The glass slides were then sealed with coverslips for microscopic observation. Microglia and astrocytes were observed and cell numbers were counted using a Neurolucida 360 system equipped with an inverted microscope (MBF Bioscience, Williston, VT, USA).

### Western blotting

Three adult UE marmosets (female, 2.8 year-old; male, 6.8 y; female, 8.8 y) were deeply anesthetized with pentobarbital after ketamine preanesthesia. After confirming the disappearance of reflexes, ice-cold PBS was transcardially perfused and subsequent cardiac arrest was confirmed. While continuing ice-cold PBS perfusion, a portion of the skull was removed and the entire brain was harvested. The gray matter of the prefrontal cortex was excised from the brain and placed in a 2-mL plastic tube, which was immediately frozen on dry ice and stored at −80°C until use. 53-93 mg of frozen prefrontal gray matter was placed in a Lysing Matrix D 2-mL tube (MP-Biomedicals, Irvine, CA, USA) cooled on ice, 800 μl of Syn-PER™ Synaptic Protein Extraction Reagent (Thermo Fisher Scientific, Waltham, MA, USA) and cOmplete mini Protease Inhibitor Cocktails (Merck, Darmstadt, Germany) was added. The tissue was homogenized by shaking the Matrix D tube at 3,200 rpm for 30 seconds using a bead beater homogenizer (μT-12, Taitec, Koshigaya, Japan). The homogenate was then centrifuged at 1200 g for 10 min. The supernatant was further centrifuged at 15,000 g for 20 min to obtain a synaptosome fraction. The synaptosome fraction was re-suspended in 700 μl of SynPER and the protein components of the synaptosome were extracted by mixing 70 μl of 100% Trichloroacetic acid and allowing it to stand for 30 min on ice. The protein extract was collected by a centrifugation at 15,000 g for 5 min. SDS-PAGE sample buffer was then added to the extract and neutralized by adding 1 M Tris. SDS-PAGE (7.5% gel, 10 or 1 μg total protein per lane, for anti-oxytocin-receptor or anti-PSD95/anti-β-actin, respectively) was performed and proteins were transferred to PVDF membranes and stained with 0.2% Ponceau-S in 5% acetic acid. The membrane was then incubated in a blocking solution containing 5% skimmed milk, 0.2% Tween20, and 0.02% NaN_3_ in PBS at 4°C, overnight. Membranes were incubated with the following primary antibodies: anti-PSD95, mouse, Novus Biologicals, #NB300-556, 1:4,000; anti-Oxytocin receptor, rabbit, Abcam, #217212, 1:100; anti-β-actin, mouse, Santa Cruz, #sc-47778, 1:4,000 in PBS-T (0.1% Tween20 in PBS) for 3.5h at room temperature. After twice of washing with PBS-T, the membranes were treated with secondary antibodies: Sheep anti-mouse IgG +HRP, Cytiva, #NIF825, 1:4,000; Goat anti-rabbit IgG +HRP, MBL, #458, 1:5,000 for 30 min at room temperature. After 4 washes with PBS-T, the blots were visualized with ECL Western Blotting Reagents, Cytiva, #RPN2109, and detected using Fusion-S solo imager (Vilber, Collégien, France).

### Statistical analysis

All data are presented as mean ± s.e.m (*n* = dendrite or axon numbers) unless otherwise stated. The Mann-Whitney rank sum test was used to analyze the data shown in Figures. 1C, 1D, 2B, 3E, 4D and Figure S1. The difference in probability distribution between cumulative plots was calculated using the Kolmogorov-Smirnov test, as shown in Figures 1E, 1F, 2C, and 4E. The differences between means were analyzed using the Friedman test, as shown in Figures 5B, 5C, Figures S3A, S3B and the Kruskal-Wallis test in Figures S6A-F. The Fisher’s exact test was used to examine the frequency of the carryover spines and that of the total spines in Figure 3C and 3D and to examine the spine survival rates for each condition in Figures 1G and 3F. The *P*-values in Figure 5J were obtained from the distribution of the simulation. The *r* and *P*-values in Figures 6A show Pearson’s correlation coefficients (*r*) and their *P*-values (*P*_r_) against zero. The concordance values (*c*) and their *P*-values (*P*_c_) in Figure 6B show the number of genes exhibiting concordant changes between the animal model and human ASD divided by the total number of genes, and the probability was calculated using the one-sided binomial test. Other statistical tests are specified in the text or figure legends or elsewhere in the **Methods** section. Quantification, simulation, and statistical analysis were performed using Microsoft Excel, GraphPad Prism 9.3.1., and Real Statistics software (for Kolmogorov-Smirnov test; www.real-statistics.com.). Analyses on spines were first conducted by an analyst blinded to the sample labels and subsequently validated by an independent analyst. In the context of axonal analysis, the quantification of fluorescence intensity was carried out in a semi-automated manner, delineating a linear region of interest (ROI) along the axon, utilizing Image-J (FIJI) software.

### Data availability

The microarray data generated in this study have been deposited at NCBI GEO under accession number GSE199560.

## Supplemental information

**Figure S1. Glial cell activation after the final imaging session was negligible.** (**A**) Activated astrocyte marker GFAP postive cell density was calculated in the fixed brain slices prepared after the final imaging session. Distinct sign of astrocyte activation under the imaging window was not detected (*P* = 0.26 at 83-115 mm cortical depth; Mann-Whitney U-test; Total 12 sites on the dorsomedial prefrontal cortex (1.1×10^-3^ mm^3^ for one site) from three marmosets). (**B**) Microglia marker Iba-1 postive cell density was calculated as in (A). Obvious sign of microglia activation under the imaging window was not detected. (*P* = 0.27 at 92-112 mm cortical depth; Mann-Whitney U-test; Total 12 sites (1.1×10^-3^ mm^3^ for one site) from three marmosets). Data are represented as mean ± s.e.m.

**Figure S2. Examination of the proximity of dendritic spine generation.** (**A**) The relationship between the clustering thresholds and differences in clustering probabilities of the newly generated spines between VPA-exposed animals and UE animals. The threshold value applied for the subsequent analyses (3 µm) was located at the lowest end of the plateau. (Related to Fig. 2). See Materials and Methods for details. (**B**) The clustered spine generation observed in the PFC of marmoset model and in the motor cortex of a mouse syndromic autism model (*MeCP2*-duplication) reported in previous literature^10^. In the graph, the clustering threshold (*T*_h_) was set to 9 µm, which was common to both mouse and marmoset data (mean ± s.e.m.; *n* = 14 and 12 dendrites in four and three UE and VPA-exposed marmosets respectively). Note that the data for mice is after four days of motor training. (**C**) The carryover spines in the marmoset PFC and those in motor cortex in the MeCP2-dup mice. (**D**) Validation of clustering bias by another Monte Carlo simulation. The new spine positions were randomly determined with a uniform distribution. This was done without pooling the spine count and dendrite length of each dendrite in this simulation, since the generation or elimination of spines may occur in specific active dendrites. Clustering probabilities for all inter-spine distances were calculated for each simulation, and the distributions of the clustering probability from 100,000 iterations are shown in (E)-(H). (**E**-**H**) Circles connected with gray lines represent probability plots of clustering events from 100,000 simulations; the actual numbers of spine clusters are represented by arrows (*P* = 0.33 and 0.0062; *n* = 23 and 65 newly generated spine pairs in 14 and 12 dendrites in UE and VPA-exposed animals, respectively) (*P* = 0.083 and 0.46; *n* = 43 and 78 newly eliminated spine pairs in 14 and 12 dendrites in UE and VPA-exposed animals, respectively). Dotted red lines show 95th percentiles. \*\*\**P* < 0.001; \*\**P* < 0.01; \**P* < 0.05; n.s., not significant.

**Figure S3. Oxytocin did not modify the proximity of spine elimination in the model marmosets.** (**A** and **B**) Mean values of 3-day total spine elimination (A) and clustered or non-clustered spine elimination(B) are shown as in Fig. 5B, C. There were no significant changes after oxytocin administration in either total spine elimination (A)(mean ± s.e.m.; *P* = 0.92, Friedman test; *n* = 12 dendrites in three VPA-exposed animals) or clustered/non-clustered spine elimination (B)(mean ± s.e.m.; *P* = 0.42 and *P* = 0.28 for clustered and non-clustered spine elimination, respectively, Friedman test; *n* = 12 dendrites in three VPA-exposed animals). (**C** and **D**) Distributions of inter-spine distances between newly generated spines (*P* > 0.85; Kolmogorov-Smirnov test; *n* = 45, 43 and 39 spine pairs for D0-3, D3-6, and D6-9, respectively) or between newly eliminated spines (*P* > 0.56; Kolmogorov-Smirnov test; *n* = 48, 38 and 48 spine pairs for D0-3, D3-6, and D6-9, respectively). n.s., not significant.

**Figure S4. Oxytocin modifies the proximity of spine generation in the model marmosets.** (**A**-**F**) Validation of clustering bias by another Monte Carlo simulation. In this simulation, the new spine positions were randomly determined with a uniform distribution without changing the spine number or length and number of each dendrite (See Fig. S2). Effects of oxytocin on clustering bias of newly generated (A-C) and eliminated (D-F) spines. The graphs are shown as in Fig. 5. (*n* = 32, 32, and 28 newly generated spine pairs and *n* = 34, 27, and 37 newly eliminated spine pairs during the D0-3, D3-6, and D6-9 periods, respectively, in three VPA-exposed animals). (**G**) The P-values expressed in logarithm from (A)-(F) are indicated. The red dotted line indicates 0.05. \*\**P* < 0.01; n.s., not significant.

**Figure S5. Oxytocin receptors are present in the synapse of the marmoset’s prefrontal cortex gray matter.** Oxytocin receptors along with post-synaptic protein PSD95 and beta-actin are detected using western blotting in the synaptosomal fraction derived from the prefrontal cortex gray matter of three adult marmosets.

**Figure S6. Oxytocin does not significantly modify axonal bouton generation or elimination.** (**A** and **B**) Mean bouton density values of ipsilateral (A) and contralateral (B) axons in VPA-exposed animals are not significantly different before and after oxytocin administration. (mean ± s.e.m.; *P* = 0.997 and *P* = 0.90 for ipsilateral and contralateral axons, respectively, Kruskal-Wallis test; *n* = 17 and 21 for ipsilateral and contralateral axons, respectively) (**C** and **D**) Three-day bouton generation (C) and elimination (D) during the D0-3, D3-6, and D6-9 periods for each axon type are shown (mean ± s.e.m.; *P* = 0.85 and *P* = 0.96 for bouton generation and elimination, respectively, Kruskal-Wallis test; *n* = 38 axons in two VPA-exposed animals). (**E** and **F**) Three-day bouton generation (E) and elimination (F) during the D0-3, D3-6, and D6-9 periods for the total axons in the VPA-exposed animals are shown as in (C) and (D) (mean ± s.e.m.; *P* = 0.85 and *P* = 0.97 for ipsilateral and contralateral bouton generation, respectively, *P* = 0.99 and *P* = 0.91 for ipsilateral and contralateral bouton elimination, respectively, Kruskal-Wallis test; *n* = 17 and 21 ipsilateral and contralateral axons in two VPA-exposed animals). n.s., not significant. (Related to Fig. 5)

## Notes

### Competing Interest Statement

The authors have declared no competing interest.

### Summary of Updates

The authors have updated the main text, including the title and abstract. Schematic drawing (Fig. 8) was removed. Supplemental files updated.

