## Supplemental information (Figure S1-S6) for "Altered projection-specific synaptic remodeling and its modification by oxytocin in an idiopathic autism marmoset model"

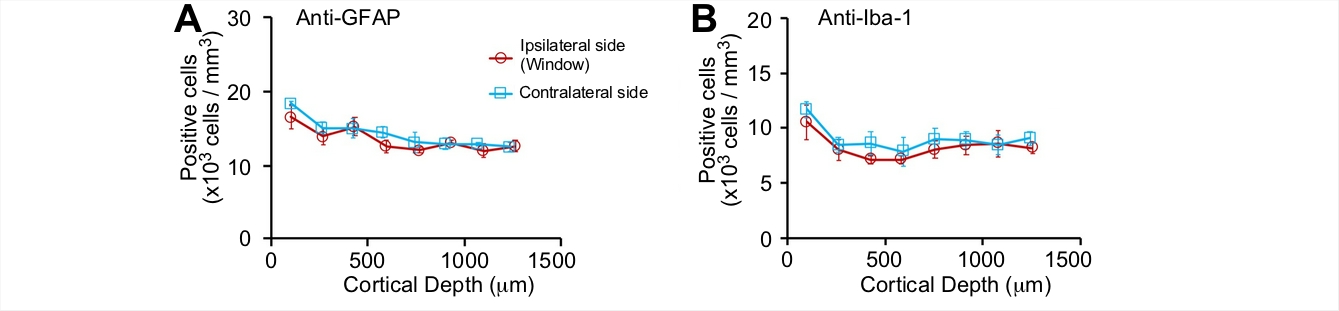

**Figure S1. Glial cell activation after the final imaging session was negligible.** (**A**) Activated astrocyte marker GFAP postive cell density was calculated in the fixed brain slices prepared after the final imaging session. Distinct sign of astrocyte activation under the imaging window was not detected (*P* = 0.26 at 83-115 mm cortical depth; Mann-Whitney U-test; Total 12 sites on the dorsomedial prefrontal cortex (1.1×10^-3^ mm^3^ for one site) from three marmosets). (**B**) Microglia marker Iba-1 postive cell density was calculated as in (A). Obvious sign of microglia activation under the imaging window was not detected. (*P* = 0.27 at 92-112 mm cortical depth; Mann-Whitney U-test; Total 12 sites (1.1×10^-3^ mm^3^ for one site) from three marmosets). Data are represented as mean ± s.e.m.

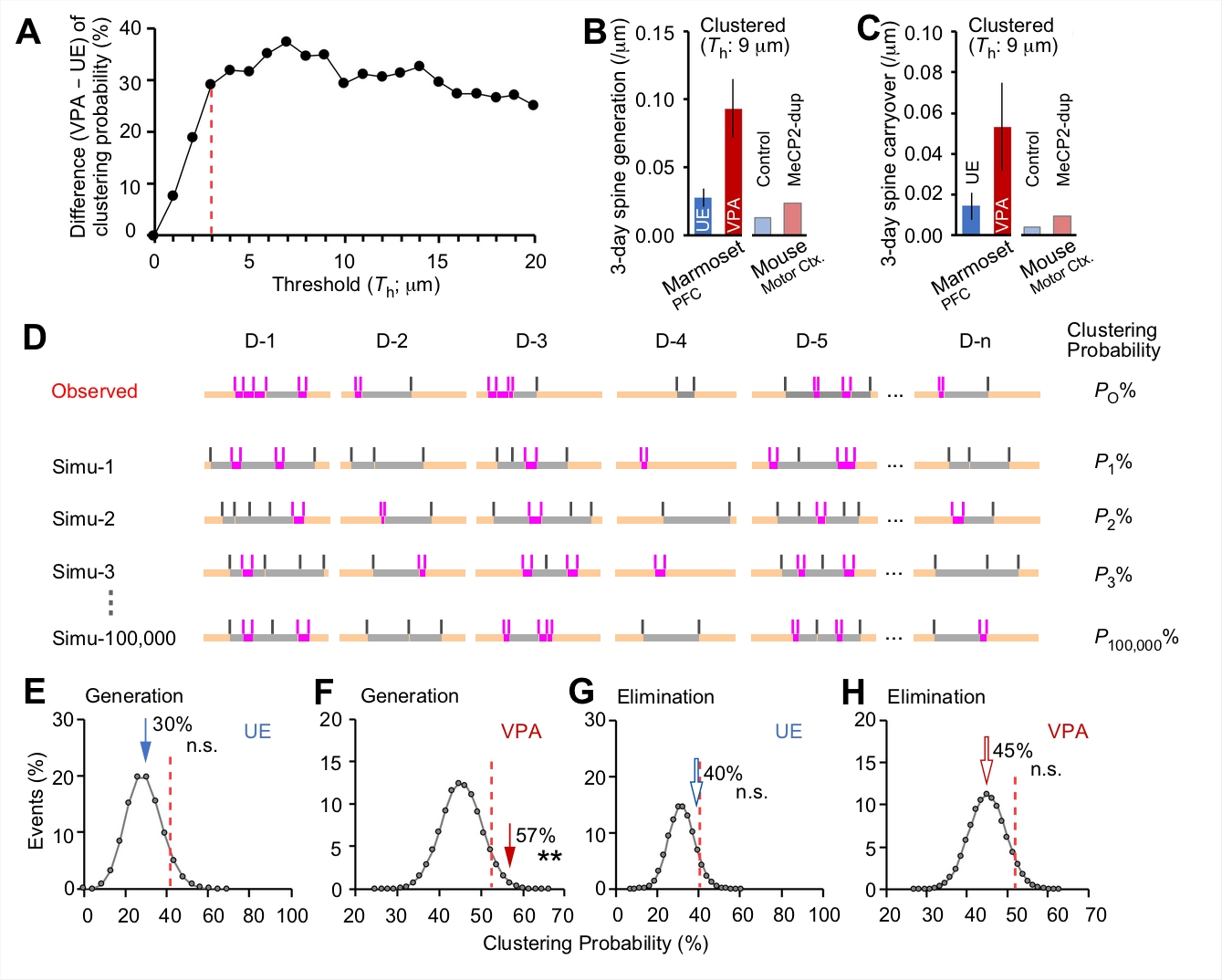

**Figure S2. Examination of the proximity of dendritic spine generation.** (**A**) The relationship between the clustering thresholds and differences in clustering probabilities of the newly generated spines between VPA-exposed animals and UE animals. The threshold value applied for the subsequent analyses (3 μm) was located at the lowest end of the plateau. (Related to Fig. 2). See STAR Methods for details. (**B**) The clustered spine generation observed in the PFC of marmoset model and in the motor cortex of a mouse syndromic autism model (*MeCP2*-duplication) reported in previous literature^10^. In the graph, the clustering threshold (*T*_h_) was set to 9 μm, which was common to both mouse and marmoset data (mean ± s.e.m.; *n* = 14 and 12 dendrites in four and three UE and VPA-exposed marmosets respectively). Note that the data for mice is after 4 days of motor training. (**C**) The carryover spines in the marmoset PFC and those in motor cortex in the MeCP2-dup mice (**D**) Validation of clustering bias by another Monte Carlo simulation. The new spine positions were randomly determined with a uniform distribution. This was done without pooling the spine count and dendrite length of each dendrite in this simulation, since the generation or elimination of spines may occur in specific active dendrites. Clustering probabilities for all inter-spine distances were calculated for each simulation, and the distributions of the clustering probability from 100,000 iterations are shown in (E)–(H). (**E**–**H**) Circles connected with gray lines represent probability plots of clustering events from 100,000 simulations; the actual numbers of spine clusters are represented by arrows (*P* = 0.33 and 0.0062; *n* = 23 and 65 newly generated spine pairs in 14 and 12 dendrites in UE and VPA-exposed animals, respectively) (*P* = 0.083 and 0.46; *n* = 43 and 78 newly eliminated spine pairs in 14 and 12 dendrites in UE and VPA-exposed animals, respectively). Dotted red lines show 95th percentiles. ****P* < 0.001; ***P* < 0.01; **P* < 0.05; n.s., not significant.

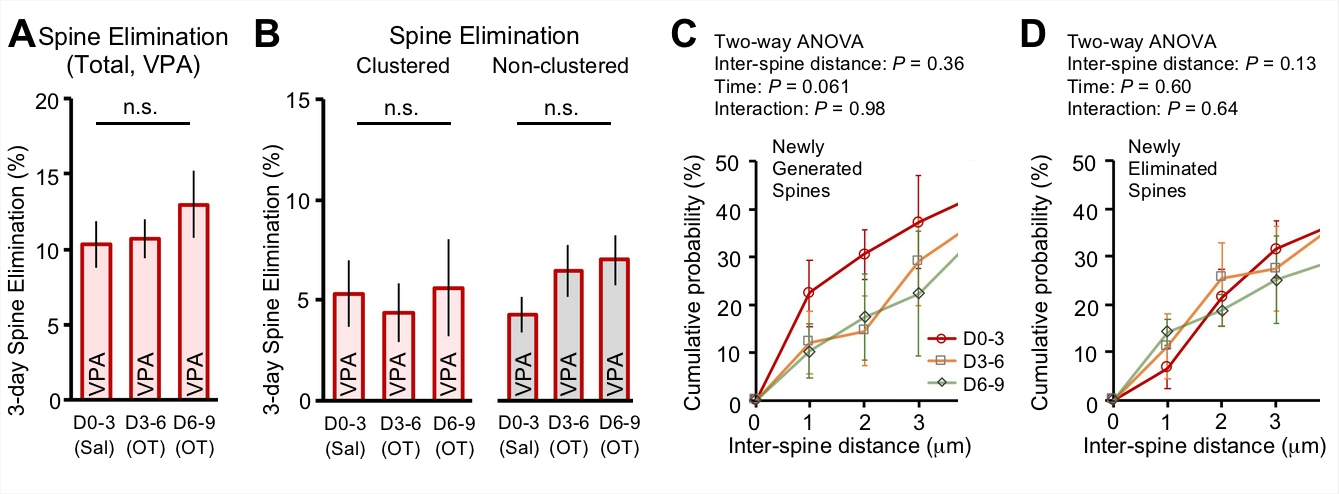

**Figure S3. Oxytocin did not modify the proximity of spine elimination in the model marmosets.** (**A** and **B**) Mean values of 3-day total spine elimination (A) and clustered or non-clustered spine elimination (B) are shown as in Fig. 5B, C. There were no significant changes after oxytocin administration in either total spine elimination (A)(mean ± s.e.m.; *P* = 0.92, Friedman test; *n* = 12 dendrites in three VPA-exposed animals) or clustered/non-clustered spine elimination (B)(mean ± s.e.m.; *P* = 0.42 and *P* = 0.28 for clustered and non-clustered spine elimination, respectively, Friedman test; *n* = 12 dendrites in three VPA-exposed animals). (**C** and **D**) Distributions of inter-spine distances between newly generated spines (*P* > 0.85; Kolmogorov-Smirnov test; *n* = 45, 43 and 39 spine pairs for D0–3, D3–6, and D6–9, respectively) or between newly eliminated spines (*P* > 0.56; Kolmogorov-Smirnov test; *n* = 48, 38 and 48 spine pairs for D0–3, D3–6, and D6–9, respectively). n.s., not significant.

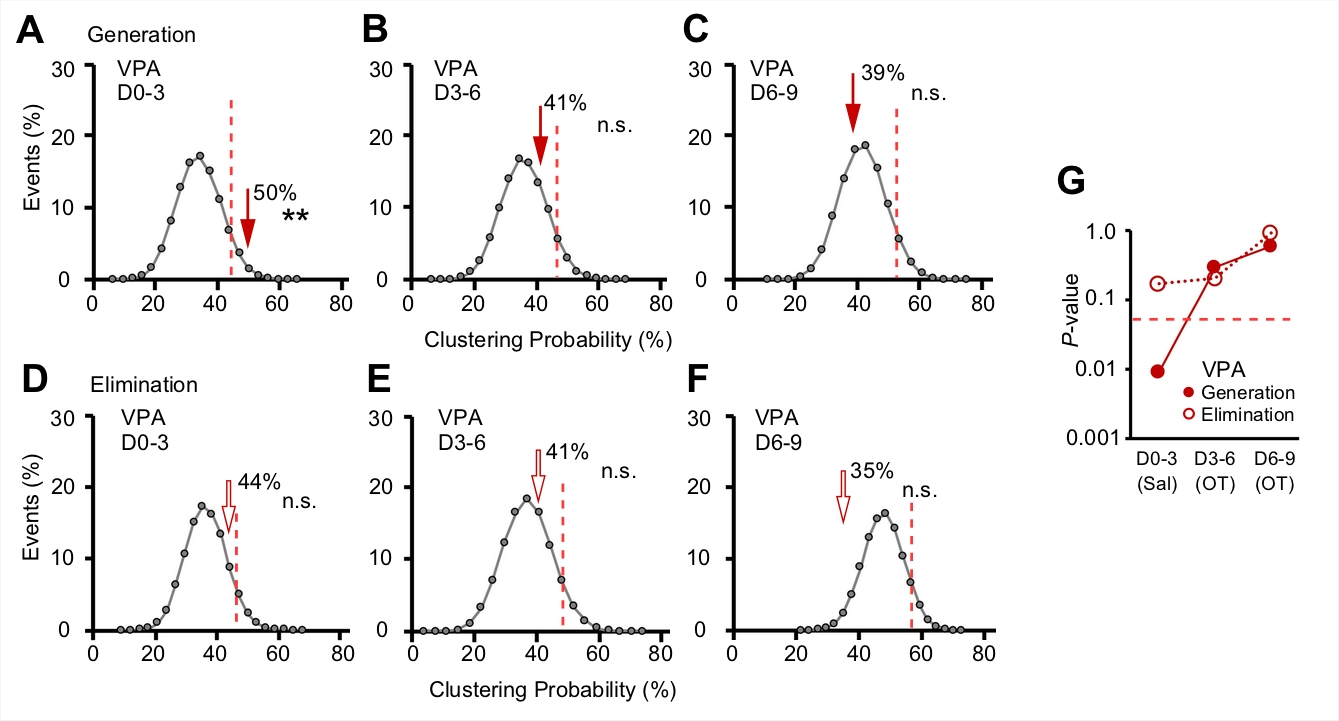

**Figure S4. Oxytocin modifies the proximity of spine generation in the model marmosets.** (**A**–**F**) Validation of clustering bias by another Monte Carlo simulation. In this simulation, the new spine positions were randomly determined with a uniform distribution without changing the spine number or length and number of each dendrite (See Fig. S2). Effects of oxytocin on clustering bias of newly generated (A–C) and eliminated (D–F) spines. The graphs are shown as in Fig. 5. (*n* = 32, 32, and 28 newly generated spine pairs and *n* = 34, 27, and 37 newly eliminated spine pairs during the D0–3, D3–6, and D6–9 periods, respectively, in three VPA-exposed animals). (**G**) The *P*-values expressed in logarithm from (A)–(F) are indicated. The red dotted line indicates 0.05. ***P* < 0.01; n.s., not significant.

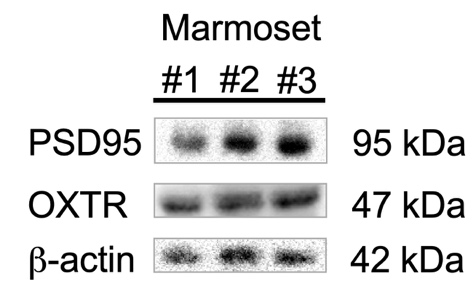

**Figure S5. Oxytocin receptors are present in the synapse of the marmoset's prefrontal cortex gray matter.** Oxytocin receptors (OXTR) along with post-synaptic protein PSD95 and β-actin are detected using western blotting in the synaptosomal fraction derived from the prefrontal cortex gray matter of three adult marmosets.

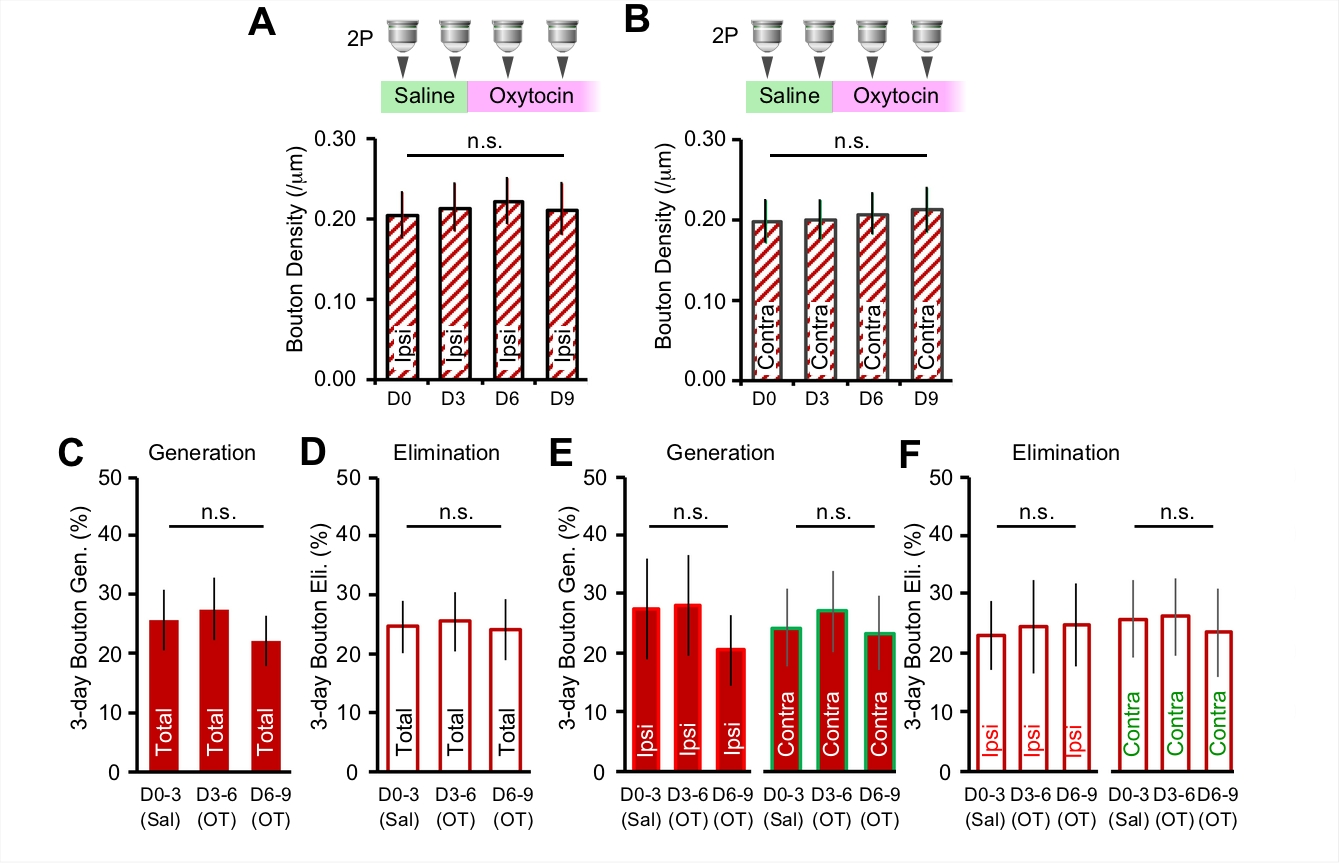

**Figure S6. Oxytocin does not significantly modify axonal bouton generation or elimination.** (**A** and **B**) Mean bouton density values of ipsilateral (A) and contralateral (B) axons in VPA-exposed animals are not significantly different before and after oxytocin administration. (mean ± s.e.m.; *P* = 0.997 and *P* = 0.90 for ipsilateral and contralateral axons, respectively, Kruskal-Wallis test; *n* = 17 and 21 for ipsilateral and contralateral axons, respectively) (**C** and **D**) Three-day bouton generation (C) and elimination (D) during the D0–3, D3–6, and D6–9 periods for each axon type are shown (mean ± s.e.m.; *P* = 0.85 and *P* = 0.96 for bouton generation and elimination, respectively, Kruskal-Wallis test; *n* = 38 axons in two VPA-exposed animals). (**E** and **F**) Three-day bouton generation (E) and elimination (F) during the D0–3, D3–6, and D6–9 periods for the total axons in the VPA-exposed animals are shown as in (C) and (D) (mean ± s.e.m.; *P* = 0.85 and *P* = 0.97 for ipsilateral and contralateral bouton generation, respectively, *P* = 0.99 and *P* = 0.91 for ipsilateral and contralateral bouton elimination, respectively, Kruskal-Wallis test; *n* = 17 and 21 ipsilateral and contralateral axons in two VPA-exposed animals). n.s., not significant. (Related to Fig. 5)
